# Kinetic modeling of ethylene biosynthesis and signaling pathways in ripening tomato fruit

**DOI:** 10.1101/2024.08.26.607967

**Authors:** Thao Minh Viet Nguyen, Dinh Thi Tran, Clara I. Mata, Bram Van de Poel, Bart M. Nicolaï, Maarten L. A. T. M. Hertog

## Abstract

- Ethylene biosynthesis and signaling are pivotal pathways in various plant aging processes, including fruit ripening. Kinetic models can be used to better understand metabolic pathways, but modeling of the ethylene-related pathways is limited and the link between these pathways remains unsolved.
- A transcriptomics-based kinetic model was developed, consisting of ordinary differential equations describing ethylene biosynthesis and signaling pathways in tomato during fruit development, ripening and post-harvest storage.
- This model was calibrated against a large volume of transcriptomic, proteomic and metabolic data during on-vine ripening of tomato fruit grown in winter and summer. The model was validated using data on off-vine postharvest ripening. The ethylene biosynthesis pathway under different conditions appeared to be largely driven by gene expression levels.
- The ethylene-regulation of fruit ripening of a heat tolerant tomato grown in different seasons is identical but with quantitative differences at the targeted omics levels. This is reflected by some of the same parameters with distinct values for summer and winter fruit. The current model is the first attempt to model the ethylene signaling pathway starting from gene expression, the various protein – protein interactions, including the link with ethylene production, internal ethylene levels and its receptors.

## Introduction

Ethylene is a key hormone regulating various plant processes including fruit ripening via a signaling pathway (McManus, 2012). The ethylene biosynthesis pathway was unraveled late 1970s (Adams & Yang, 1979; Yu *et al*., 1979). It starts from the conversion of S-adenosylmethionine (SAM) to 1-aminocyclopropane-1-carboxylic acid (ACC) catalyzed by ACC synthase (ACS). In tomato, SAM is a common substrate involved in three kind of pathways: ethylene biosynthesis, biosynthesis of polyamines, and transmethylation reactions (Van de Poel *et al*., 2013). In tomato, there are 14 *ACS* genes, *ACS1A* and *ACS1B* to *ACS13* (http://tomexpress.toulouse.inra.fr/). During tomato ripening, the transcripts of *ACS1A, ACS2, ACS4* and *ACS6* are strongly expressed, while *ACS3, ACS5, ACS7* and *ACS8* are undetectable (Nakatsuka *et al*., 1998; Barry *et al*., 2000; Van de Poel *et al*., 2012). The second reaction of ethylene biosynthesis entails the conversion of ACC to ethylene by ACC oxidase (ACO) (Hamilton *et al*., 1990). ACC can also be converted to 1-(malonylamino)cyclopropane-1-carboxylic acid (MACC) (Amrhein *et al*., 1981; Martin & Saftner, 1995). Besides MACC formation, ACC can be also conjugated into γ-glutamyl-ACC (GACC) (Martin *et al*., 1995), and jasmonyl-ACC (JA-ACC) (Staswick & Tiryaki, 2004). While ACS is well-known as a rate-limiting enzyme, ACO can also be rate-limiting under certain conditions (Dorling & McManus, 2012; Van de Poel & Van Der Straeten, 2014). Seven ACO members, ACO1 to ACO7, have been identified in tomato (Houben & Van de Poel, 2019), four of which (*ACO1, ACO3, ACO5* and *ACO6*) are differentially expressed during fruit ripening (Barry *et al*., 1996, 2000; Nakatsuka *et al*., 1998).

In the late 1980s, studies on the triple response, an ethylene-specific response of dark-grown seedlings, were key in isolating the first ethylene receptor ETR1 in *Arabidopsis*, opening up new insights into ethylene reception and signal transduction (Bleecker *et al*., 1988; Roman *et al*., 1995; Chen & Bleecker, 1995). Subsequent research identified a multi-gene family of seven ethylene receptor (ETR) members in tomato (Lin *et al*., 2009b). Mutant studies showed that these receptors are negative regulators (Ciardi *et al*., 2000; Tieman *et al*., 2000). These ETRs can form homo- and heterodimers which are stabilized through covalent disulfide bonds at their N-terminal domain (Schaller *et al*., 1995; Gao *et al*., 2008; Chen *et al*., 2010; Kamiyoshihara *et al*., 2022). The second component of ethylene signaling is the constitutive triple response (CTR) protein, also a negative regulator (Lin *et al*., 2008a; Liu *et al*., 2015). Studies on ethylene signaling in *Arabidopsis* reported that CTR1 localizes in the endoplasmic reticulum (ER) (Clark *et al*., 1998; Gao *et al*., 2003), yet Park *et al*. (2023) found CTR1 in both ER and nucleus. Four CTR homologs have been identified in tomato (Klee & Giovannoni, 2011). In the absence of ethylene, ETRs signal to the CTRs which phosphorylate and inhibit the ethylene-insensitive protein 2 (EIN2), a positive regulator, consequently preventing ethylene signaling cascades (Ju *et al*., 2012). On ethylene binding, ETR protein degradation is triggered. As the signal is no longer transmitted to the CTRs, EIN2 is being activated, allowing the cleavage of EIN2 into C-terminal (EIN2_Cterm_) and N-terminal (EIN2_Nterm_) domains. While the function of EIN2_Nterm_ remains unclear, EIN2_Cterm_ has two roles in ethylene signaling. EIN2_Cterm_ binds *EIN3-binding F-box 1* (*EBF1*) and *EBF2* mRNA to direct them to cytoplasmic processing bodies, resulting in their degradation, thus preventing the ubiquitination degradation of EIN3 and EIN3 like (EIL) (Li *et al*., 2015). EIN2_Cterm_ is also transported to the nucleus to activate ethylene responses via the EIN3/EIL family and ethylene response factors (Zhang *et al*., 2017). Full-length EIN2 was also found in the nucleus, proposing an alternative signaling in which EIN2 translocates to the nucleus to cleave the C-terminal domain (Zhang *et al*., 2020). However, the translocation of an integral membrane protein to the nucleus remains enigmatic.

Kinetic modelling is a quantitative mechanistic approach to untangle complex biological networks and their regulation (Rios-Estepa & Lange, 2007; Schallau & Junker, 2010). Still, kinetic models of hormone-related biological pathways in plant are limited. Génard and Gouble (2005) developed a model of ethylene production by peach fruit involving the effects of temperature, O_2_, and CO_2_. Van de Poel *et al*. (2014) developed a transcriptomic-based kinetic model providing an integrated overview of ethylene biosynthesis during fruit development and ripening. However, due to a lack of experimental data on protein abundance, the link between transcriptomics and proteomics remained implicit. Solely based on gene expression data, ACO1 was assumed to be the main contributor to ACO activity controlling ethylene production (Van de Poel *et al*., 2014a). This assumption was premature because work from Nguyen et al. (2023) found that also ACO3, ACO5 and ACO6 are important in regulating ACO activity.

So far, no attempt was made to develop a detailed model on ethylene signaling. Gwanpua et al., (2017) modelled the impact of 1-Methylcyclopropene (1-MCP) on apple ripening based on a lumped description of ethylene biosynthesis and signaling. However, such model cannot describe the specific contribution of individual receptor members, the role of homodimers and heterodimers, the contribution of CTRs and EIN2, downstream signaling components, and the feedback to ethylene biosynthesis. While our understanding of plant ethylene biosynthesis, its regulation, and the ethylene signal transmission has been consolidated (Wang *et al*., 2002; Chen *et al*., 2020), these pathways were mostly studied in isolation. The current study presents a comprehensive integrated kinetic model on ethylene biosynthesis and signaling during tomato fruit ripening and postharvest storage covering its main transcript, proteins and metabolites.

## Materials and Methods

### Plant materials and biochemical characteristics

Experimental data (on-vine data) published by Nguyen et al. (2023) were used to calibrate the current model. The model was validated against additional experimental data on off-vine ripening tomato fruit using fruit of the same cultivar grown in the same seasons. Fruit in were grown in the same field (21°09’07.9”N 106°00’59.9”E) harvested in winter (18 – 24 °C) and summer (21 – 32 °C) seasons. For the on-vine dataset, winter and summer fruit were harvested at immature green (IMG), mature green (MG), breaker (BR), turning (TRN), orange (ORG), light red (LR) and red ripe (RR) stages. Additionally, harvested RR tomatoes were stored for 6 d and 12 d at 18 °C and 80 % RH. Regarding the off-vine dataset, MG winter and summer tomato fruit were harvested and either treated with 5 μL L^-1^ 1-Methylcyclopropene (1-MCP), 100 μL L^-1^ ethylene for 24 h at 18 °C or exposed to ambient atmosphere (non-treated control fruit). All fruit were stored at 18 °C and 80 % RH. For both seasons the control fruit were sampled for further analyses when they reached BR, ORG, LR, and RR stages, and two post-climacteric stages (RR+6, RR+12). 1-MCP and ethylene treated tomatoes were sampled at the same time as the control fruit regardless their actual ripening stage. For 1-MCP treated fruit, an extra sampling stage at RR+18 was introduced. Fruit color (Hue angle) and ethylene production rate were measured at each sampling time following Nguyen *et al*. (2023). Immediately afterwards, pericarp tissue was flash frozen in liquid nitrogen, ground and stored at -80 °C for the analyses of ethylene biosynthesis and signaling related transcripts (*ACO1*–*7, ACS1A* and *B, ACS2*–8, *ETR1*– *7, CTR1*–*3*, and *EIN2*), corresponding proteins, and substrates (ACC and MACC). Measurements of ACC and MACC content (μmol kg^-1^), ACO and ACS activity (nmol kg^-1^ s^-1^), gene expression (/) and protein abundance (μmol kg^-1^ protein) related to ethylene biosynthesis and signaling were identical to Nguyen *et al*. (2023). The measurements of protein abundance were only conducted for the on-vine dataset.

### Transformation of categorical maturity stage into a continuous time scale

Tomato maturity stages are typically defined using discrete color stages. Nguyen et al. (2023) harvested the fruit simultaneously at different color stages, so no information is available on the time needed to progress from one stage to the next. However, to model the time changes in ethylene biosynthesis and signaling a continuous time scale is required. To conduct this transformation, the sampling days of off-vine validation data were used as a reference (see Supplementary table 1) as this dataset involved the same cultivar grown under the identical condition, sharing the same experimental sampling protocol except that fruit were all harvested at the MG stage and then allowed to ripen. We assumed the time intervals between maturity stages to be the same for tomatoes ripened on-vine. Since the on-vine experiment started from the immature green (IMG) stage, the time interval between immature green (IMG) and MG was missing. Based on previous experience we assumed this to be one day.

### Model development, from concept to math

#### Conceptual model

Our model was developed based on the current understanding of ethylene biosynthesis and the canonical ethylene signaling pathway. From literature we extracted a conceptual model (Fig. 1) which was used to formulate a kinetic ordinary differential equation (ODE) based model. Gene expression data were used as model inputs while the outputs are protein abundance, enzyme activity, concentrations of intermediates and ethylene production rate. Mass action based kinetic rates were assumed. Fig. 1 visualizes the two pathways with their related variables and corresponding rate constants. Reaction rates are denoted with the symbol *k*_x,y_, with ‘x’ the reaction type and ‘y’ the name of the enzyme or another identifier of the reaction. Reaction types and subscripts are listed in Table 1. A detailed list of symbols is provided in Supplementary table 2.

**Table 1.**
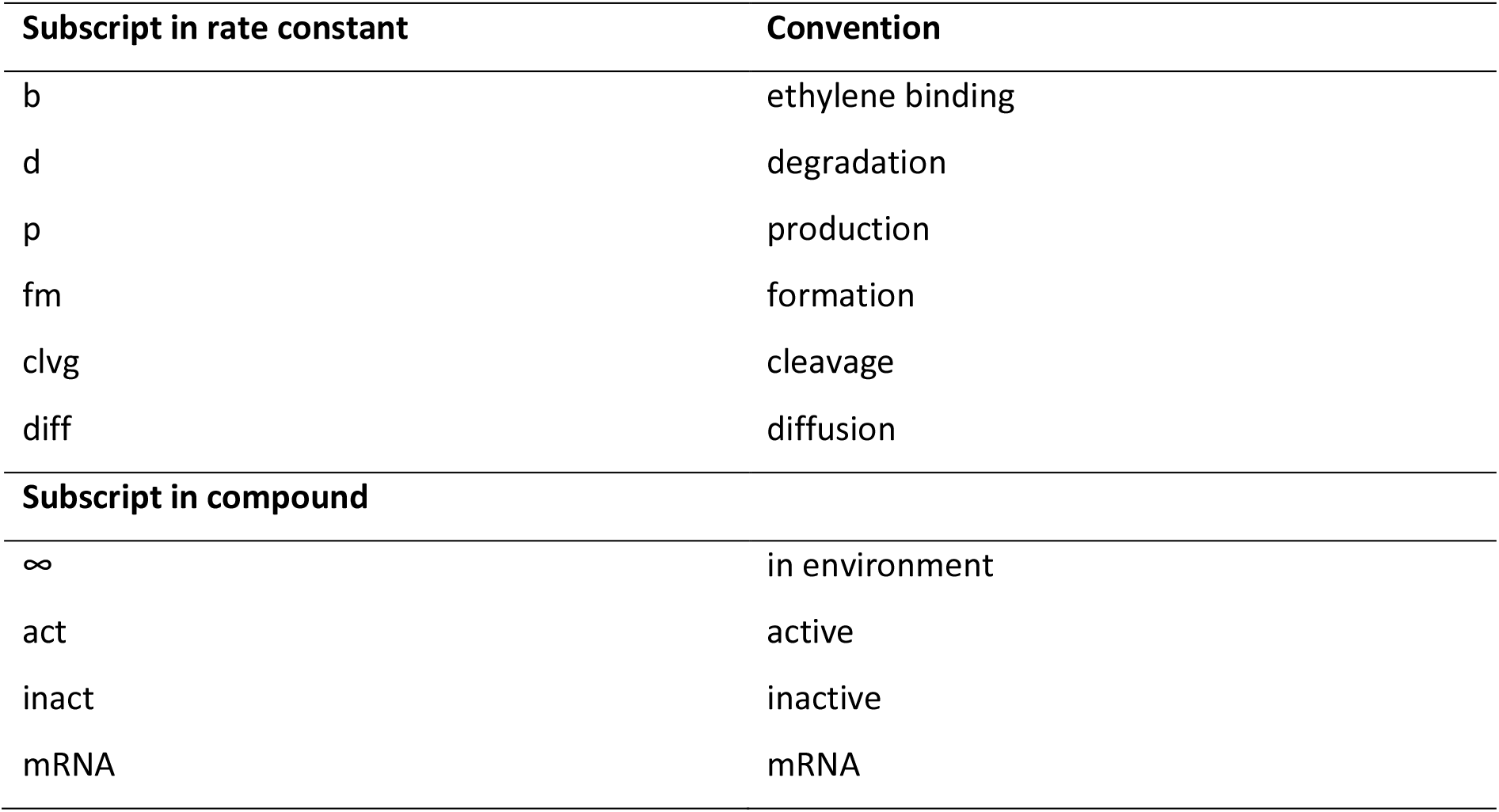
The subscript convention in rate constant and in the compound.

**Fig. 1.**
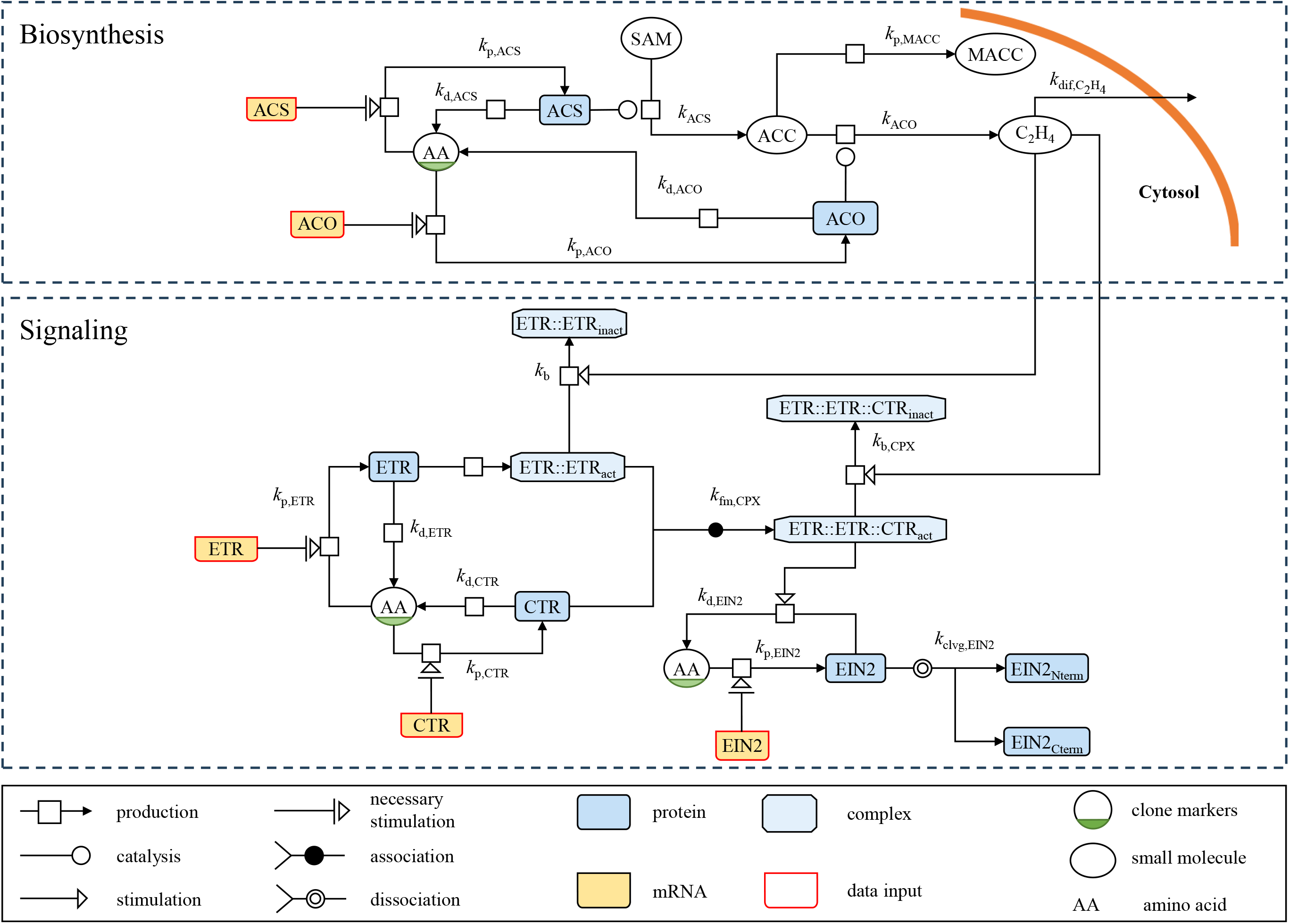
Simplified scheme of ethylene biosynthesis and signaling pathway as implemented in the kinetic model. The precursor of ethylene, ACC, is first produced from SAM catalyzed by ACS members with a rate constant *k*_ACS_. ACC is then converted to internal ethylene (C_2_H_4_) catalyzed by ACO members with rate constants *k*_ACO_. ACC is also converted to MACC with a rate constant *k*_p,MACC_. A portion of internal ethylene diffuses out of the tissue at a certain diffusion rate 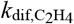. Specific ACO members are translated from their corresponding mRNAs by specific rate constants *k*_p,ACO_. The total of ACS members is translated from their corresponding mRNAs with a generic rate constant *k*_p,ACS_. ACO and ACS members are degraded with a generic rate constant *k*_d,ACO_ and *k*_d,ACS_, respectively. For ethylene signaling, CTR and ETR members are translated from their total corresponding mRNAs with rate constants *k*_p,CTR_ and *k*_p,ETR_ respectively. The degradation of CTR and ETR members is represented by the rate constants *k*_d,CTR_ and *k*_d,ETR_. ETR members interact with each other, forming homo/heterodimer ETR :: ETR_act_, subsequently interacting with CTR to form the complex ETR :: ETR :: CTR_act_ with a rate constant *k*_fm,CPX_. The binding of ethylene deactivates ETR :: ETR :: CTR_act_ to ETR :: ETR :: CTR_inact_, with a rate constant *k*_b,CPX_. The binding of ethylene deactivates ETR :: ETR_act_, with the rate constants *k*_b_. The translation and degradation of EIN2 is expressed by the rate constant *k*_p,EIN2_ and *k*_d,EIN2_, respectively. The later step is stimulated by the complex ETR :: ETR :: CTR_act_. The cleavage of EIN2 into EIN2_Cterm_ is represented by the rate constants *k*_clvg,EIN2_. ETR refers to all ETR members in subfamily I and II. ETR :: ETR_act_ refers to active homo- or heterodimers. ETR :: ETR :: CTR_act_, and ETR :: ETR :: CTR_inact_ refer to active homo- or heterodimer/ CTR complexes and inactive homo- or heterodimer/ CTR complexes in general. The input data are the gene expression. The outputs include protein abundance, enzymatic activities, metabolic contents, and ethylene production rate. The putative SAM concentration profile was predicted during calibration. All reactions in this model were assumed to be irreversible reactions. The graphical notation is according to Rougny et al., (2019).

#### Ethylene biosynthesis: from mRNA to ethylene

The kinetic model describing ethylene biosynthesis was modified from Van de Poel et al., (2014). In their model, ACO1 was assumed to be the main contributor to ACO activity, and thus ethylene production. However, the protein results from Nguyen et al. (2023) revealed that ACO3, ACO5 and ACO6 also play an important role in ethylene biosynthesis. For ACS members, it was reported that the transcripts of *ACS2, ACS4*, and *ACS6* were highly expressed throughout fruit development, ripening, and postharvest storage (Van de Poel *et al*., 2012; Nguyen *et al*., 2023). Therefore, this study considered all of them (*ACO1, ACO3, ACO5, ACO6, ACS2, ACS4*, and *ACS6)*. To account for the difference in house-keeping gene expression observed between winter and summer fruit (Nguyen et al., 2023), a correction factor *f*_HKG_ was employed. Since the mRNA transcript levels were used as inputs for the translation steps, post-transcriptional modifications are implicitly included in our model. In the ethylene biosynthesis submodel, all post-translational modifications negatively regulating protein turnover were combined into a single protein degradation step (*k*_d,ACO_, *k*_d,ACS_). The turnover of each ACO member was assumed to result from a specific translation rate *k*_p,ACO*i*_ and a generic degradation rate *k*_d,ACO_ (Eq. 1).

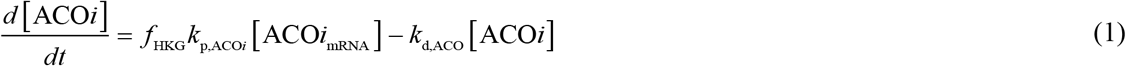

with *i* ∈{1,3,5, 6}.

Since there was no information on ACS abundance, assigning a single parameter for the translation of each ACS member results in an over-parameterized model. Therefore, the overall ACS protein turnover is calculated as the net balance between a general translation (*k*_p,ACS_) and degradation (*k*_d,ACS_) term (Eq. 2).

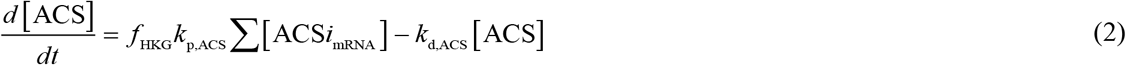

for *i* ∈{2, 4,6}.

The production of ACC, MACC and ethylene is highly related to SAM content (Bürstenbinder & Sauter, 2012; Van de Poel *et al*., 2012) which was shown to be time dependent. Unfortunately, no SAM data was available. Hence, we reconstructed a putative SAM concentration time profile using a piecewise cubic Hermite polynomial with six equidistantly spaced positive knots requiring the SAM concentration at these six knots to be estimated as model parameters.

While three ACC derivatives are known (MACC, GACC and JA-ACC), Peiser and Fa Yang, (1998) showed MACC to be the major conjugate of ACC in ripening tomato. Therefore, we ignored GACC and JA-ACC assuming ACC content is the net balance between its synthesis by ACS, ethylene production by ACO, and conversion to MACC (Eqs. (3), (4)). The ethylene accumulation results from its synthesis from ACC and its exchange with the environment (Eq. 5).

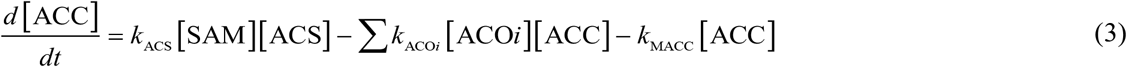

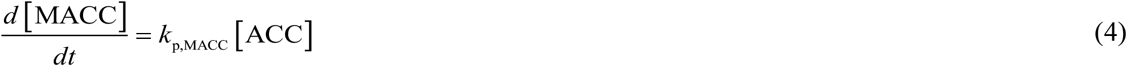

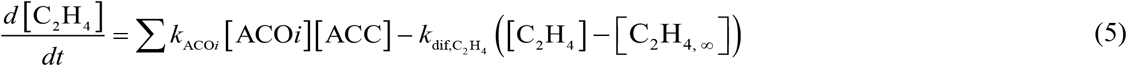

#### Ethylene signal transmission

In this study, the kinetic model is expanded with the ethylene signaling pathway. *ETR1, ETR2, ETR3, ETR4, ETR6, ETR7, CTR1* are the main genes in ethylene signal transmission (Kevany *et al*., 2007; Mata *et al*., 2018; Nguyen *et al*., 2023), which were used as key transcripts in this model. In tomato, ETR1, ETR2 and ETR3 belong to subfamily I, whereas ETR4, ETR5, ETR6, and ETR7 belong to subfamily II (Chen *et al*., 2018). Kamiyoshihara et al. (2022) observed that ETR in subfamily I interact with themselves and with other members to form homodimer and heterodimers, while ETR in subfamily II did not. Their study also showed that ETR from subfamily I interacts with ETR from subfamily II. Subsequently, all types of dimers interact with CTRs (CTR1 in our study) and activate them (Clark *et al*., 1998; Huang *et al*., 2003). This complex mechanism is modelled in a simplified way, immediately forming the active homo- or heterodimer/CTR1 complex (ETR::ETR::CTR1_act_). One homo- or heterodimer molecule was assumed to interact with one CTR1 molecule. In the presence of ethylene, the binding of ethylene results in conformational receptor changes, that reduce the interaction and activation of CTR1 (Binder, 2020). There is no information whether the detached CTR1 is reactivated. However, it was reported that the protein levels of ETR::ETR::CTR1_act_ complexes decreased at higher ethylene concentration (Shakeel *et al*., 2015). Therefore, we assumed that the binding of ethylene deactivates the complex ETR::ETR::CTR1_act_, including CTR1. Ethylene binds to ETR dimers (Schaller & Bleecker, 1995; O’Malley *et al*., 2005), inactivating and degrading them (Kevany *et al*., 2007; Chen *et al*., 2007). The ubiquitin-related degradation of ETR members and CTR1 is referred to as general degradation. Taken together, the turnover of each free ETR (monomer) is the net balance between a specific translation (*k*_p,ETR*i*_, *k*_p,ETR*j*_), general degradation (*k*_d,ETR_), the complex formation with CTR1 (*k*_fm,CPX_), and the inactivation of dimers by ethylene 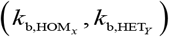 (Eqs. (6), (7)). The difference between these two equations is in the dimer formation of ETRs from subfamily I and II.

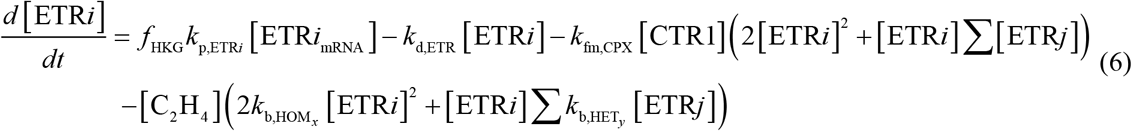

with 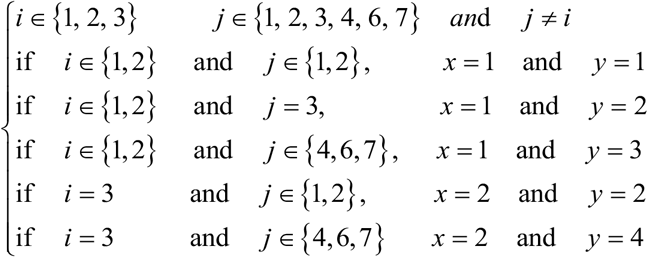

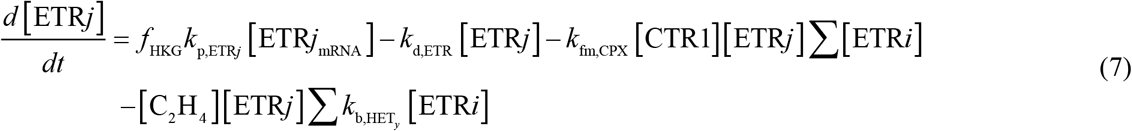

with 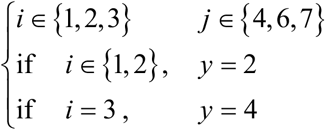

The turnover of CTR1 is the net balance between the translation (*k*_p,CTR1_), degradation (*k*_d,CTR1_), and the complex formation with receptor dimers (*k*_p,CPX_) (Eq. 8).

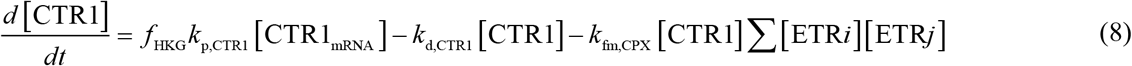

with *i* ∈{1, 2, 3} *j* ∈{1, 2, 3, 4, 6, 7} *j* ≥ *i*

The turnover of each ETR::ETR::CTR1_act_ equals the complex formation, general degradation, and degradation enhanced by ethylene (Eq. 9).

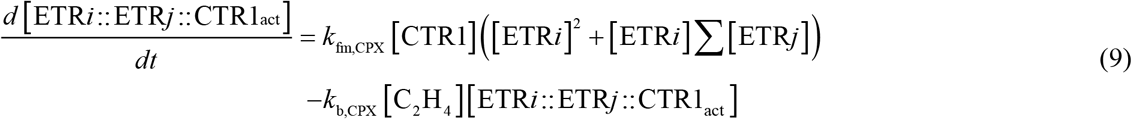

with *i* ∈{1, 2, 3} *j* ∈{1, 2, 3, 4, 6, 7} *i* ≠ *j* and *j* ≥ *i*

In the absence of ethylene, ETR::ETR::CTR1_act_ complexes inhibit EIN2 by phosphorylation, which triggers the ubiquitin-related degradation (Ju *et al*., 2012; Nguyen *et al*., 2013). In the presence of ethylene, ETR::ETR::CTR1_act_ becomes inactivated, reducing phosphorylation of EIN2. There is no information on the specific affinity of the different ETR::ETR::CTR1_act_ complexes to EIN2. Therefore, we assume the degradation of EIN2 to depend on the total abundance of ETR::ETR::CTR1_act_. EIN2 cleaves into two parts, the C-terminal part (EIN2_Cterm_) and N-terminal part. The EIN2 turnover is the net balance between translation (*k*_p,EIN2_), degradation (*k*_d,EIN2_) enhanced by ETR :: ETR :: CTR1_act_, and cleavage into EIN2_Cterm_ (*k*_clvg,EIN2_) resulting in Eq. 10. This model does not include the possible role of the full-length EIN2 in the nucleus.

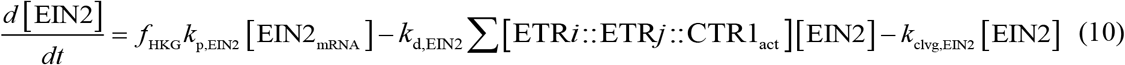

with *i* ∈{1, 2, 3} *j* ∈{1, 2, 3, 4, 6, 7} *j* ≥ *i*

#### Output relations

Multiple output relations were defined to link the ODE variables to the measured variables. The ethylene production rate by the intact fruit was assumed to be proportional to the ethylene levels inside the fruit. The *in vitro* activity of ACO and ACS was assumed to reflect the *in vivo* activities through some conversion factors *c*_ACO_ and *c*_ACS_. The modelled concentrations of the various receptors were collated into their total abundance combining free proteins with proteins bound in the ETR::ETR::CTR1_act_ complex as the LC-MS based protocols were not able to discriminate between the different configurations. The inactive dimers (ETR :: ETR_inact_) and the inactive complexes (ETR :: ETR ::CTR1_inact_) were assumed to rapidly degrade, thereby no longer contributing to the measured levels of ETR or CTR1. Since CTR1 was measured in a membrane protein extract, it represents the amount of CTR1 interacting with receptors and, therefore, represents the total abundance of ETR::ETR::CTR1_act_ complexes.

#### Model implementation

The model was implemented in its ordinary differential equation (ODE) form, and the kinetic parameters were estimated using OptiPa (version 6.3p; Hertog *et al*., 2007). For model calibration we used the datasets on winter and summer tomato fruit ripened on-vine. During model calibration, we first estimated most parameters as being in common between the two seasons, gradually turning parameters to be seasonally specific to improve the model fit. Gene expression data were used as inputs and the outputs were protein abundance, enzyme activity, concentrations of intermediates and ethylene production rate. We used weighted least-squares optimization with the Levenberg-Marquardt method for parameter estimation. A standard normal transformation was applied to each measurement variable. The full model script is provided in Supplementary Dataset 1.

To validate the model, Monte-Carlo simulations were performed to predict the off-vine data including prediction uncertainties. Starting from the calibration data, confidence intervals were established for the main model parameters using 150 bootstrapping runs. To correct for heteroscedasticity in the calibration data, a Box-Cox transformation was applied. For each bootstrap dataset, parameters were re-estimated resulting in a population of 150 parameter sets. From these, parameters’ distributions and their covariance structure were obtained. These were used as an input to the Monte-Carlo analysis to simulate the off-vine data. Seven experimental datasets on off-vine ripening were used, three for winter and four for summer crop. Both seasons contained data on control fruit, 1-MCP and ethylene treated fruit. The control experiment in summer was done in duplicate but for the current purpose combined into one. The validation data contained no information on SAM, therefore the putative SAM concentration profiles were re-estimated. As there was no information on MACCT gene expression, the rate constant *k*_MACC_ was re-estimated as well. The initial values of ACC, MACC, and ethylene production rate were fixed at their measured values. The initial abundances of the proteins were set at the simulated values for the MG stage from the calibration results. All remaining parameters were inherited from the calibration data. Since the on-vine experiments used for model calibration started at the IMG stage, while the off-vine experiments started at the MG stage, the time interval was corrected by subtracting one day (the assumed time interval between the IMG and MG stage). The Monte-Carlo simulations for the off-vine datasets thus relied largely on the bootstrapped calibration parameters combined with the measured gene expressions and initial values of the validation experiments and a few re-estimated parameters motivated by the absence of experimental data.

## Results

### Model calibration

Most parameters were estimated from calibration data on ethylene production rate, the concentrations of ethylene biosynthesis intermediates, the abundance and the activity of ethylene biosynthesis enzymes, and the abundance of signaling proteins obtained from on-vine experiments. During model calibration, we sequentially optimized parameters for ethylene biosynthesis followed by those for ethylene signaling. Eventhough the ethylene biosynthesis model was partially based on Van de Poel et al. (2014a), given the modifications made and the different cultivar involved, all parameter values needed to be re-estimated for the current data. As the information in the data was sometimes limited or largely implicit, some parameters showed high correlations. For pairs of highly correlated parameters, one was arbitrarily fixed, resulting in a set of fixed and estimated parameters (see table 2 and 3). All fixed parameters were used in common for winter and summer fruit, except for *f*_HKG_. The HKGs gene expression of winter fruit was taken as a reference, using *f*_HKG_ =1. Based on the average difference in HKGs expression between winter and summer fruit, the *f*_HKG_ for summer fruit was set to 2.07. The measured protein abundances related to ethylene biosynthesis and signaling were expressed per unit of total protein (TP) and membrane protein (MP), respectively. However, the model output was expressed per unit of tissue mass. Therefore, two conversion factors, *c*_TP_ and *c*_MP_, were introduced (Supplementary Dataset 1).

**Table 2.** Fixed parameter values for ethylene biosynthesis and signaling applied during model calibration on the on-vine data.

| Fixed parameters | Value | Unit |
| --- | --- | --- |
| $k_{\text{d},\text{ACS}}$ | 0.1 | $\text{d}^{-1}$ |
| $k_{\text{ACS}}$ | $8 \cdot 10^5$ | $\text{kg mol}^{-1} \text{d}^{-1}$ |
| $k_{\text{ACO3}}$ | $8 \cdot 10^5$ | $\text{kg mol}^{-1} \text{d}^{-1}$ |
| $f_{\text{HKG, winter}}$ | 1 | / |
| $f_{\text{HKG, summer}}$ | 2.07 | / |
| $k_{d,ETR}$ | 0.1 | $d^{-1}$ |
| $k_{d,CTR1}$ | 0 | $d^{-1}$ |
| $k_{d,EIN2}$ | 0.02 | $d^{-1}$ |
| $k_{fm,CPX}$ | $2 \cdot 10^{12}$ | $kg^2 mol^{-2} d^{-1}$ |
| $k_b,CPX$ | 0.1 | $d^{-1}$ |
| $k_{clvg,EIN2}$ | 0.1 | $d^{-1}$ |

**Table 3.** Estimated parameter values for ethylene biosynthesis and signaling obtained from the model calibration on the on-vine data.

| Parameter estimates | Season |  | Unit |
| --- | --- | --- | --- |
|  | Winter tomato | Summer tomato |  |
| | Value ± SD | Value ± SD | |
| $k_{p,ACO1}$ | $213 \pm 37$ | $111 \pm 11$ | $\mu mol kg^{-1} d^{-1}$ |
| $k_{p,ACO3}$ | $0.267 \pm 0.076$ | $0.00513 \pm 0.02260$ | $\mu mol kg^{-1} d^{-1}$ |
| $k_{p,ACO5}$ | $0.443 \pm 0.043$ | $0.407 \pm 0.087$ | $\mu mol kg^{-1} d^{-1}$ |
| $k_{p,ACO6}$ | $0.199 \pm 0.028$ | $0.303 \pm 0.039$ | $\mu mol kg^{-1} d^{-1}$ |
| $k_{d,ACO}$ | $0.784 \pm 0.007$ | $0.263 \pm 0.014$ | $d^{-1}$ |
| $k_{p,ACS}$ | $1.22 \pm 0.06$ | $103 \pm 13.4$ | $\mu mol kg^{-1} d^{-1}$ |
| $k_{ACO1}$ | $0.796 \pm 1.61$ | $8.90 \pm 3.54$ | $kg mol^{-1} d^{-1}$ |
| $k_{ACO5}$ | $4.54 \cdot 10^5 \pm 1.34 \cdot 10^5$ | $0.098 \pm 0.154$ | $kg mol^{-1} d^{-1}$ |
| $k_{ACO6}$ | $1.59 \cdot 10^2 \pm 6.30 \cdot 10^3$ | $1.36 \cdot 10^4 \pm 1.63 \cdot 10^4$ | $kg mol^{-1} d^{-1}$ |
| $k_{MACC}$ | $1.14 \pm 0.177$ | $0.796 \pm 0.071$ | $d^{-1}$ |
| $k_{p,ETR1}$ | $91.0 \pm 30.7$ | $591 \pm 756$ | $\mu mol kg^{-1} d^{-1}$ |
| $k_{p,ETR2}$ | $0.326 \pm 0.651$ | $337 \pm 538$ | $\mu mol kg^{-1} d^{-1}$ |
| $k_{p,ETR3}$ | $1.77 \pm 1.36$ | $2.81 \pm 1.21$ | $\mu mol kg^{-1} d^{-1}$ |
| $k_{p,ETR4}$ | $5.81 \cdot 10^{-3} \pm 1.04 \cdot 10^{-3}$ | $2.04 \cdot 10^{-3} \pm 2.87 \cdot 10^{-4}$ | $\mu mol kg^{-1} d^{-1}$ |
| $k_{p,ETR6}$ | $0.503 \pm 0.080$ | $0.727 \pm 0.095$ | $\mu mol kg^{-1} d^{-1}$ |
| $k_{p,ETR7}$ | $2.01 \pm 0.46$ | $0.680 \pm 0.172$ | $\mu mol kg^{-1} d^{-1}$ |
| $k_{p,CTR1}$ | $0.0118 \pm 0.0012$ | $0.0247 \pm 0.004$ | $\mu mol kg^{-1} d^{-1}$ |
| $k_{p,EIN2}$ | $0.127 \pm 0.014$ | $0.0797 \pm 0.0048$ | $\mu mol kg^{-1} d^{-1}$ |
| $k_{b,HOM1}$ | $1.68 \cdot 10^{25} \pm 1.06 \cdot 10^{25}$ | $2.14 \cdot 10^{26} \pm 3.35 \cdot 10^{26}$ | $kg^2 mol^{-2} d^{-1}$ |
| $k_{b,HET1}$ | $5.32 \cdot 10^{24} \pm 1.10 \cdot 10^{25}$ | $2.59 \cdot 10^{28} \pm 3.27 \cdot 10^{28}$ | $kg^2 mol^{-2} d^{-1}$ |
| $k_{b,HET3}$ | $3.91 \cdot 10^{24} \pm 8.46 \cdot 10^{23}$ | $1.09 \cdot 10^{23} \pm 6.6 \cdot 10^{22}$ | $kg^2 mol^{-2} d^{-1}$ |
| $k_{b,HOM2}$ | $1.07 \cdot 10^{23} \pm 3.54 \cdot 10^{23}$ | $5.98 \cdot 10^{22} \pm 3.44 \cdot 10^{22}$ | $kg^2 mol^{-2} d^{-1}$ |
| $k_{b,HET2}$ | $8.83 \cdot 10^{24} \pm 7.39 \cdot 10^{24}$ | $2.82 \cdot 10^{24} \pm 2.05 \cdot 10^{24}$ | $kg^2 mol^{-2} d^{-1}$ |
| $k_{b,HET4}$ | $3.21 \cdot 10^{22} \pm 1.14 \cdot 10^{21}$ | $0.99 \cdot 10^{20} \pm 3.34 \cdot 10^{21}$ | $kg^2 mol^{-2} d^{-1}$ |

| Parameters estimated in common for both winter and summer fruit |  |  |
| --- | --- | --- |
| $k_{\text{dif,C}_2\text{H}_4}$ | $1.17 \pm 0.00150$ | $\text{d}^{-1}$ |
| $C_{\text{ACO}}$ | $1290 \pm 196$ | / |
| $C_{\text{ACS}}$ | $16.3 \pm 1.24$ | / |

The initial concentration of ACC, MACC, C_2_H_4_, ACO1, ACO5, ACO6, ETR1, ETR2, ETR3, ETR4, ETR6, ETR7, CTR1 and EIN2 were fixed based on the measured starting values (Supplementary table 3). The initial protein abundance of ACO3 and ACS was arbitrarily fixed based on their gene expression and the relationship between gene expression and the protein abundance of ACO1, ACO3, ACO6. The initial concentration of each ETR::ETR::CTR1_act_ was fixed such that for any specific ETR, the sum of all its involved complexes was not higher than the initial measured abundance of that receptor. In this way the total abundance of every ETR monomer, including those incorporated in complexes with CTR1, was always smaller than its measured initial abundance.

Few model parameters were estimated in common between winter and summer fruit (Table 3). The conversion factors between *in vivo* and *in vitro* activity (*c*_ACO_, *c*_ACS_), and 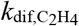 were estimated in common for both seasons as they were measured under identical lab conditions. As gas diffusion rate in tissue depend on the tissue’s micro stucture (Ho *et al*., 2010; Xiao *et al*., 2023), but given we used the same cultivar for both seasons, the diffusion rate 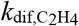 was asssumed similar and estimated in common.

The remaining rates of translation, and degradation, and the rates of enzyme activity were estimated separately for winter and summer fruit as the aim was to study the impact of growing season. Finally, the degradation rates of all proteins of the same family were assumed to be similar as there was insufficient experimental information about protein specific degradation.

### Model performance using on-vine ripening data

The kinetic model predictions, using the measured gene expression data (Supplementary Fig. 1) as input together with the parameter values from Table 2 and 3 and the initial values from Supplementary table 3, are presented in Fig. 2-4 together with the measurements. In general, the model fits the experimental data on ethylene biosynthesis for both seasons well The putative SAM concentration profile was estimated to optimally match the observed model outputs in terms of ACC, MACC and ethylene production (Supplementary Fig. 2). The high initial concentration of SAM, coupled with its steep decrease during early stages of ripening, ensures adequate substrate for ACC and subsequently MACC and ethylene. In summer tomato, the higher initial concentration of SAM together with its steeper decrease was associated with a more rapid increase in ACC, MACC, and ethylene production rate. The ACC content was predicted well except for a slight underestimation during storage of summer fruit (Fig. 2a). The simulation of MACC matched the data except for the overestimation during the late stage of ripening (Fig. 2b). For both winter and summer fruit, the *in vitro* activities of ACO and ACS and the ethylene production rate were well predicted (Fig. 2c-e).

**Fig. 2.**
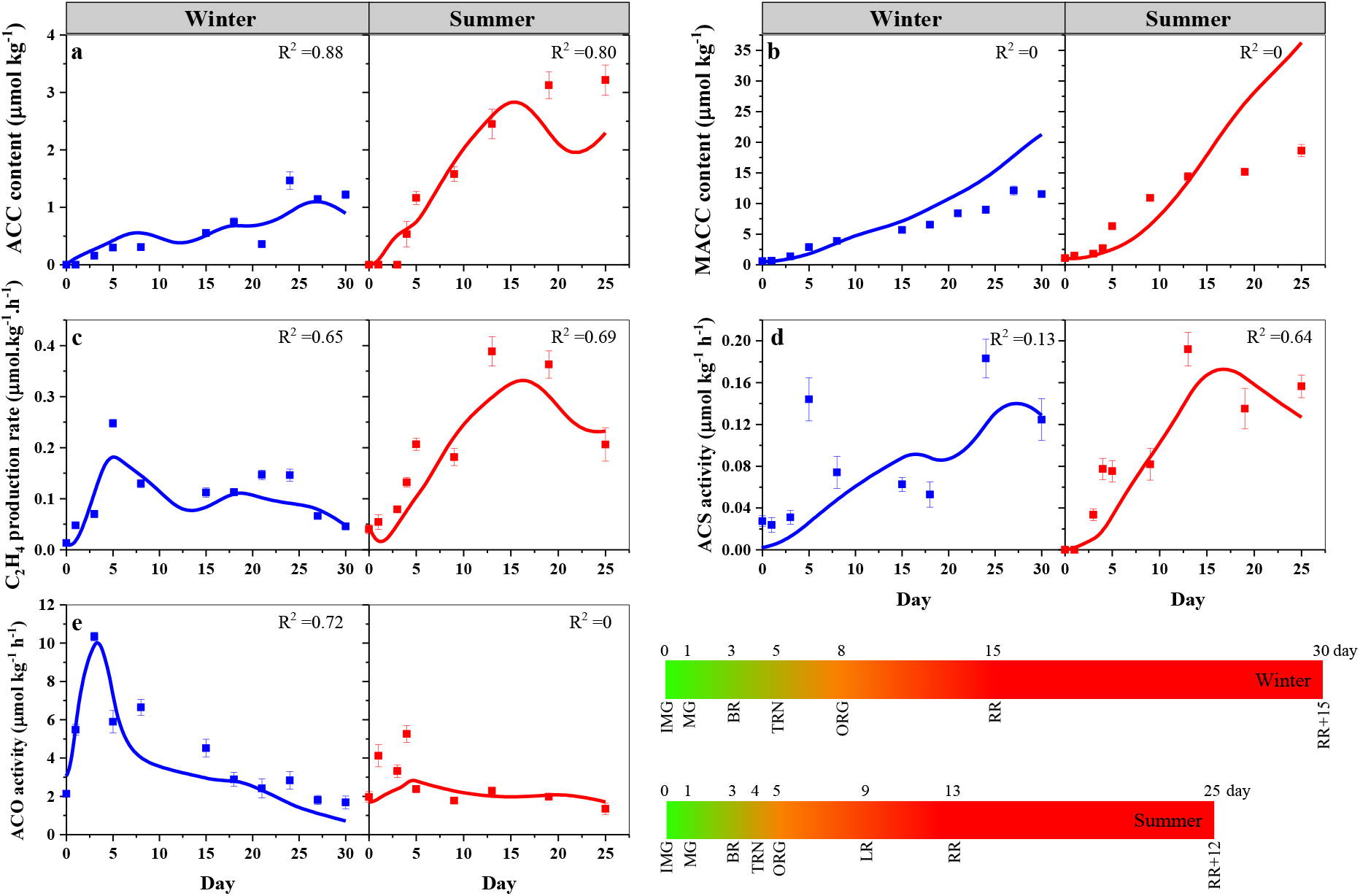
Model results of the ethylene biosynthesis pathway of winter and summer tomatoes during fruit ripening and postharvest storage. (a) ACS activity, (b) ACC content, (c) ACO activity, (d) Ethylene production rate, (e) MACC content. The lines are the model predictions, while the points are measurement data. Error bars represent the standard error of the mean (n = 5 for winter fruit and n = 4 for summer fruit).

**Fig. 3.**
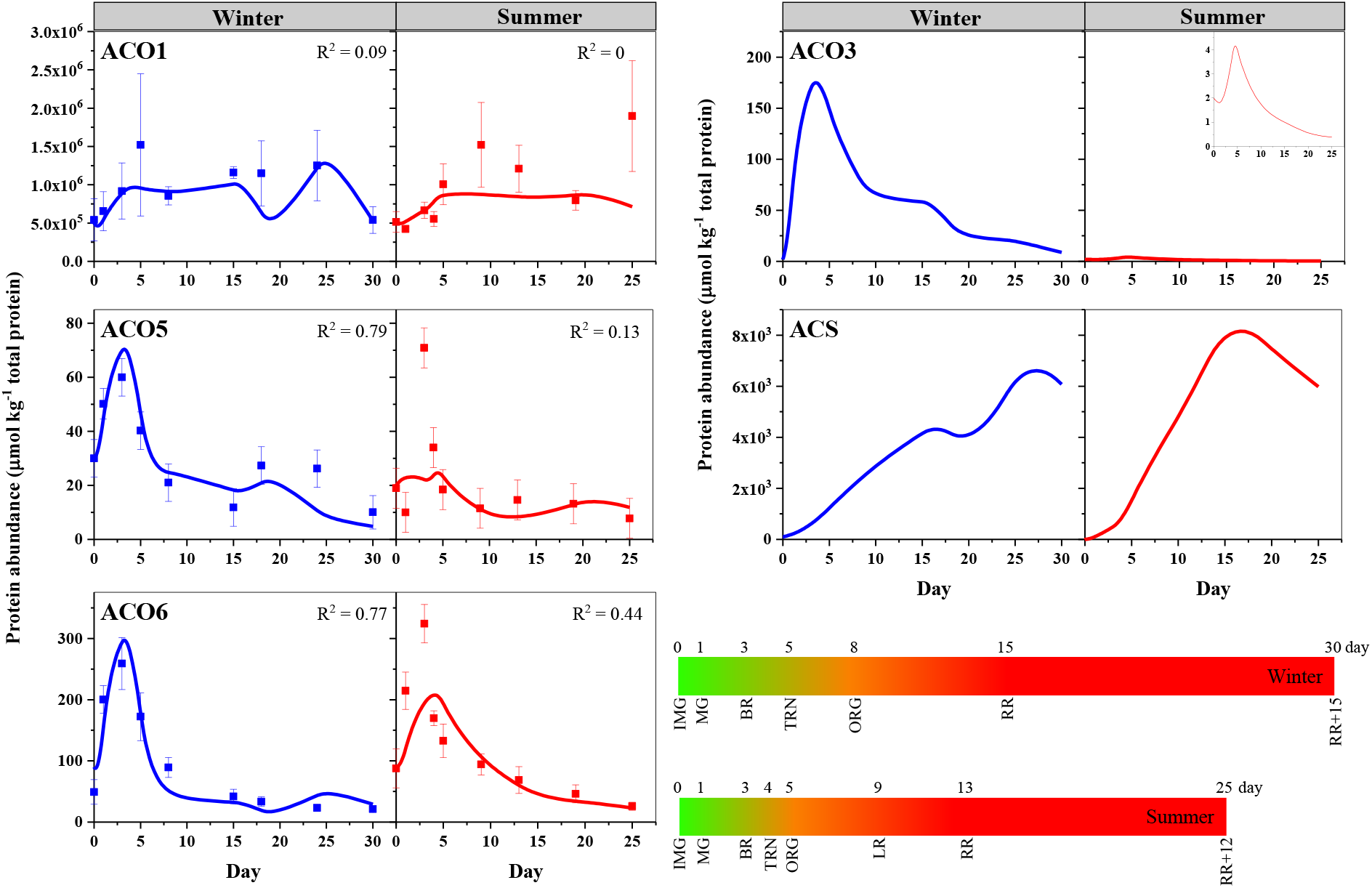
Model results of ACOs and ACS protein abundance of winter and summer tomatoes during fruit ripening. The lines are the model predictions, while the points are measurement data. Error bars represent the standard error of the mean (n = 5 for winter fruit and n = 4 for summer fruit).

**Fig. 4.**
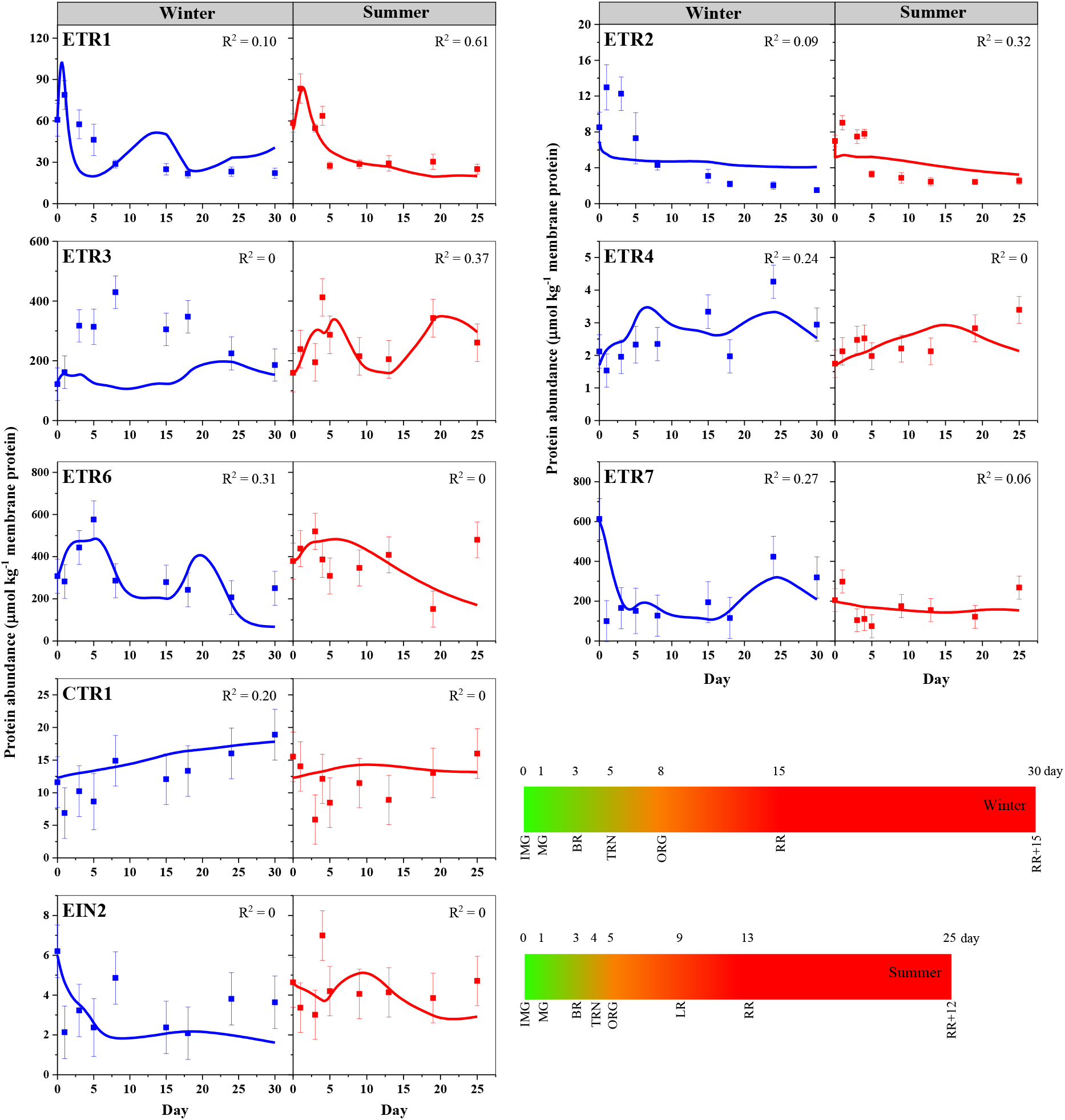
Results of signaling protein abundance of winter and summer tomatoes during fruit ripening. The lines are the model predictions, while the points are measurement data. Error bars represent the standard error of the mean (n = 5 for winter fruit and n = 4 for summer fruit).

The model well explained the protein abundance of individual ACO members (Fig. 3). In both winter and summer tomatoes, the low value of *k*_ACO1_ was compensated by the high abundance of ACO1 as compared to the other ACO isozymes (Table 3). Overall in winter fruit, ACO1 and ACO6 only marginally contribute to ACO activity, while ACO5, and to a lesser extent ACO3, are the main drivers of ACO activity and thus ethylene production (Fig. 2 and Supplementary Fig. 3). On the other hand, in summer fruit, ACO1 contributes most to the overall ACO acitivity. Since there was no experimental data on ACO3, *k*_ACO3_ was fixed, and the protein level of ACO3 was estimated. This estimation was used to indirectly predict ACO3 contribution to the ACO activity and to ethylene production (Fig. 3). The simulation showed the protein abundance of ACO3 in winter tomato to be much higher than in summer fruit.

The signaling protein abundance data matched the predictions well except for ETR2 and ETR3 during winter fruit ripening (Fig. 4). The ethylene-enhanced degradation of ETR1 and ETR2 homo- and heterodimers was assumed to proceed with the same rate constant 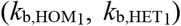. Likewise, the ethylene-enhanced degradation of heterodimers between ETR1 and ETR2 and receptors in subfamily II was characterized by the same parameter 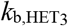. The ethylene-enhanced degradation of homo- and heterodimers containing ETR3 each had their own rate constant (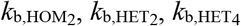; Table 3). Due to insufficient information about specific dimers, the related parameters were unidentifiable as indicated by their large confidence intervals (Table 3). Different approaches were applied during model calibration but those uncertain estimates, together with the lack-of-fit of ETR2 and underestimation of ETR3 could not be resolved. The model explained the time course of CTR1 and EIN2 abundance during both winter and summer rather well.

### Model validation using off-vine ripening data

The calibrated model was validated using off-vine data of the same tomato cultivar ‘Savior’ grown during the same winter and summer season either or not treated with ethylene or 1-methylcyclopropene (1-MCP). The measured gene expression data were used as model input (Supplementary Fig. 4). The gene expression data of the treated fruit implicitly included the impact of the treatments. The putative SAM profile was re-estimated for each experimental condition (Supplementary Fig. 5) showing in all cases a clear decline during the early period of ripening. There was no difference in putative SAM concentration between control and ethylene treated winter/summer tomatoes as both shared a similar behavior in terms of ACC, MACC, and ethylene production. In contrast, 1-MCP treated winter/summer tomatoes showed a putative SAM pattern different from the control, which aligned with the delayed accumulation of ACC, MACC and ethylene production rate. The rate constants and initial values used are summarized in Supplementary table 4. Note that the time axis was expressed relative to color stages in both on- and off-vine experiments. Therefore, the results do not provide information on a possible difference in rate of ripening between winter or summer or between on-vine and off-vine.

The Monte-Carlo based predictions for ethylene biosynthesis are shown in Fig. 5, 6, and 7. In general, ethylene biosynthesis was largely explained by the underlying gene expression. Since there is no significant difference in gene expression between non-treated and ethylene treated tomatoes (Supplementary Fig. 4), the results of their model prediction were similar (compare Fig. 5a-e to Fig. 6a-e). The observed inhibition of ethylene biosynthesis by 1-MCP matched the predictions (Fig. 7a-e). During the early stages of ripening, the ACC content appeared to be overestimated in the validation experiments, except for the 1-MCP treated fruit (Fig. 5a, 6a, 7a). Meanwhile, the ACC content in both non-treated, ethylene treated, and 1-MCP treated summer tomato during the later stages of ripening was underestimated. Furthermore, the model could not well explain the ACS activity (Fig. 5d, 6d, 7d).

**Fig. 5.**
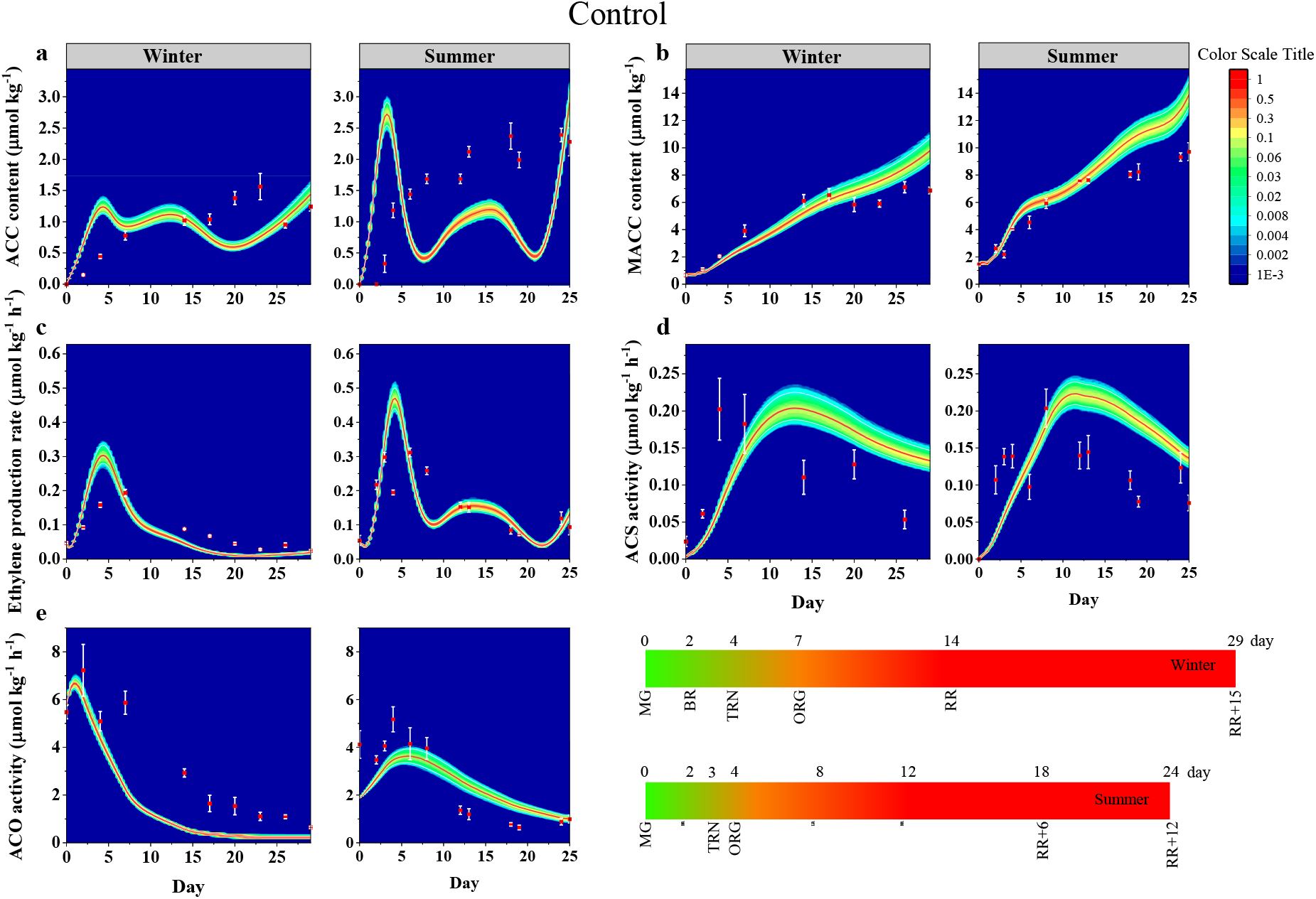
Prediction of the ethylene biosynthesis of control winter and summer tomatoes during off-vine postharvest ripening. (a) ACC content; (b) MACC content; (c) ethylene production rate; (d) ACS activity; (e) ACO activity. Heat plots represent the distribution results from a Monte Carlo simulation. The red and white lines represent the mean and 95 % of confident interval, respectively. The points are measurement data. Error bars represent the standard error of the mean (n = 5 for winter fruit and n = 4 for summer fruit).

**Fig. 6.**
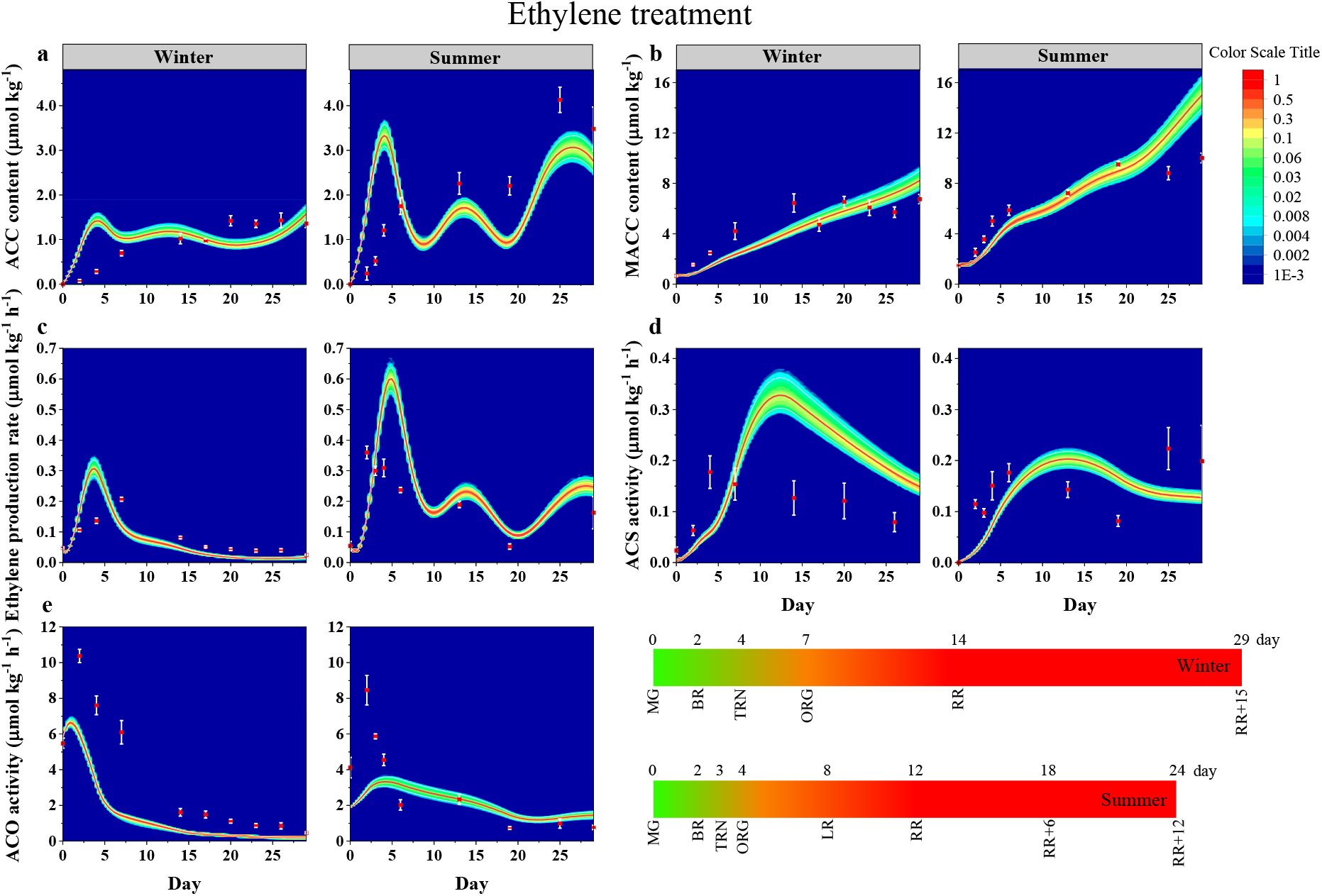
Prediction of ethylene biosynthesis of ethylene treated winter and summer tomatoes during off-vine postharvest ripening. (a) ACC content; (b) MACC content; (c) ethylene production rate; (d) ACS activity; (e) ACO activity. Heat plots represent the distribution results from a Monte Carlo simulation. The red and white lines represent the mean and 95 % of confident interval, respectively. The points are measurement data. Error bars represent the standard error of the mean (n = 5 for winter fruit and n = 4 for summer fruit).

**Fig. 7.**
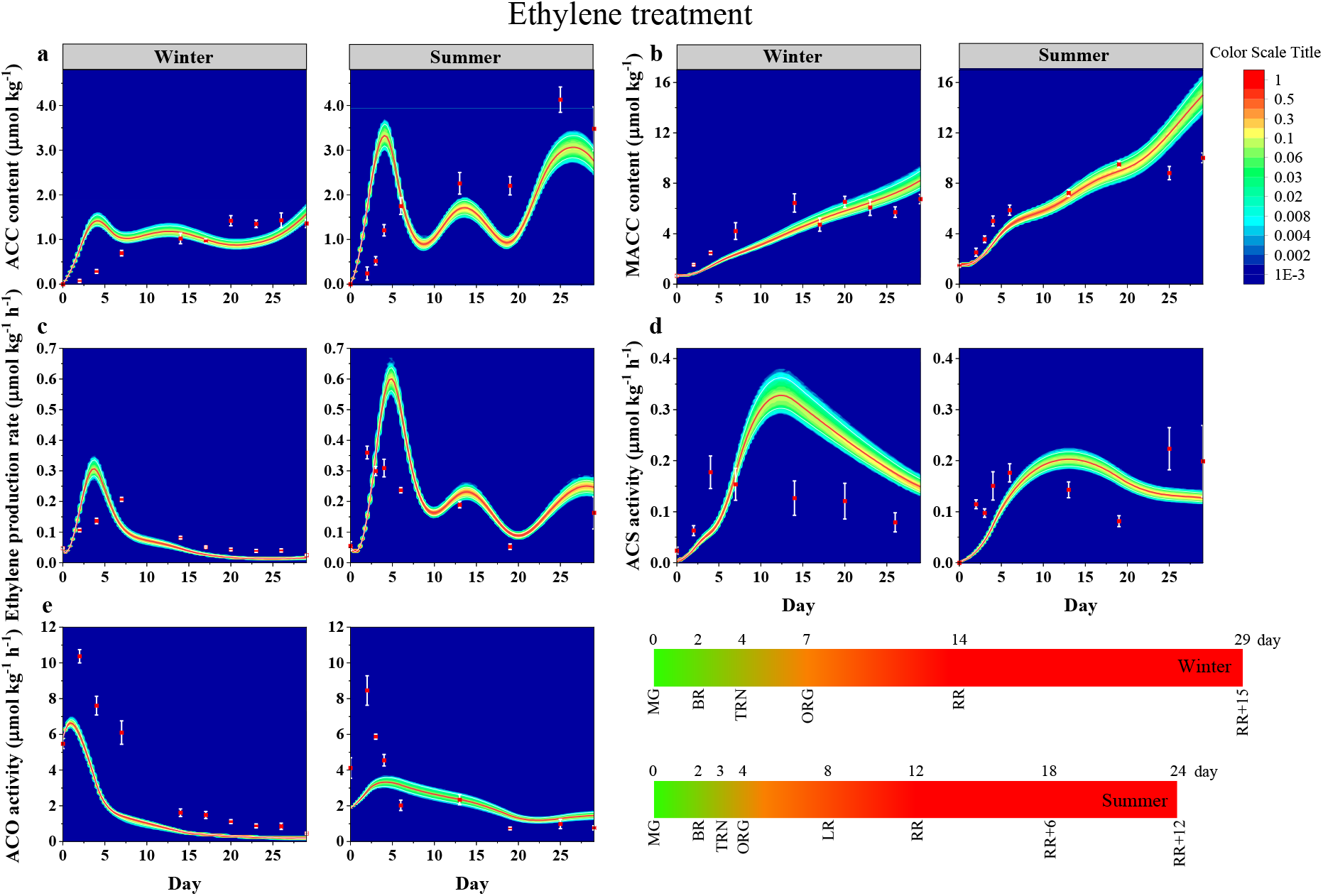
Prediction of the ethylene biosynthesis of 1-MCP treated winter and summer tomatoes during off-vine postharvest ripening. (a) ACC content; (b) MACC content; (c) ethylene production rate; (d) ACS activity; (e) ACO activity. Heat plots represent the distribution results from a Monte Carlo simulation. The red and white lines represent the mean and 95 % of confident interval, respectively. The points are measurement data. Error bars represent the standard error of the mean (n = 5 for winter fruit and n = 4 for summer fruit).

Since no experimental data on protein abundance were available, the model predicts the patterns of ACO and ACS isozymes (Supplementary Fig. 6-8) and signaling proteins (Supplementary Figure 9-11) but cannot be challenged by any experimental data. A low level of ACO3 abundance was forecasted for all summer tomatoes. Additionally, the model indicates that 1-MCP suppresses the translation of ACO members, and ACO3 in particular, during the earlier stages of ripening (Supplementary Fig. 8). Among the signaling proteins, the estimated ETR3 abundance in summer tomato was lower than that in winter fruit (Supplementary Fig. 9-11). In addition, 1-MCP was predicted to strongly affect the abundance of most receptors (Supplementary Fig. 11).

## Discussion

### Assumptions underlying some model parameters

Our previous study showed an apparent difference in gene expression between winter and summer fruit due to the altered expression of HKGs (Nguyen *et al*., 2023) which is captured by *f*_HKG_. Thanks to adding this term in the model we can correct for the lack of constant HKG still allowing proper interpretation of the experimental data. Ideally, other HKG whose expression is relatively stable during high temperature should be identified making this correction term redundant (Karkute *et al*., 2021). The degradation rate constants of ACS, ETRs and EIN2 were fixed with a small values (0.1 d^-1^) based on previous studies (Van de Poel *et al*., 2014a; Belouah *et al*., 2019). The rate constant of ETR::ETR::CTR1_act_ complex formation *k*_p,CPX_ was set at a high value assuming all CTR1 immediately interacts with ETRs. Since CTR1 abundance was much lower than any ETR abundance, together with the previous assumption, the degradation of free CTR1 was assumed negligible. Ethylene binds to CTR-free receptor dimers with different affinities (Schaller *et al*., 1995; O’Malley *et al*., 2005), while little is known about its binding affinity to specific ETR::ETR::CTR1_act_ complexes. Therefore, we assumed these to be identical. The EIN2 degradation parameter *k*_d,EIN2_ was kept at a low value to take into account the contribution of the ETR::ETR::CTR1_act_ complex to its degradation. The EIN2 degradation and EIN2 cleavage reaction are competitive. Giving a high value for the EIN2 cleavage rate constant might result in a high amount of EIN2_Cterm_ production in the earlier stages of ripening. However, only the abundance of membrane-bound full-length EIN2 was measured, excluding EIN2 C-term. To take into account the role of EIN2 degradation, it was assumed that 10 % of EIN2 cleaved into EIN2_Cterm_.

### The ethylene biosynthesis model is transferable to different tomato cultivar

The submodel on ethylene biosynthesis was based on Van de Poel *et al*. (2014a). After parameter estimation, the calibration data on ethylene biosynthesis was well explained, indicating that the model structure is transfereable to other cultivars. Given the rate 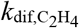, the ethylene diffusivity of cv. ‘Savior’ used in this study, is lower than that of cv. ‘Bonaparte’, used by Van de Poel et al., (2014a). This will be reflecting cultivar difference in fruit (micro)structure as also observed in pomefruit (Ho *et al*., 2010).

### Modeling ethylene biosynthesis: insights and limitations

Valuable insights can be obtained from the model starting with the putative SAM profile being different from that reported for cv. ‘Bonaparte’ (Van de Poel *et al*., 2013) (Supplementary Fig. 2). This may be due to the fact that the putative SAM profile was solely based on the ethylene biosynthesis, ignoring other pathways (e.g. biosynthesis of polyamines and transmethylation reactions) in which it is involved (Van de Poel *et al*., 2013). The underestimated ACC content might be indicative of additional ACC transport between tissues (Van de Poel *et al*., 2014b) and cell compartments (Guy & Kende, 1984; Saftner & Martin, 1993) occuring differently in winter and summer fruit. The overerestimation in MACC content during fruit ripening could be due to the lack of data on ACC-N-malonyl transferase (MACCT) or the synthesis of other ACC conjugates (GACC and JA-ACC). Besides the ACC concentration, the production of MACC is also regulated by MACCT which is indirectly controlled by its gene expression (Van de Poel *et al*., 2014a). Currently *MACCT* levels were assumed to be constant resulting in a time independent rate constant *k*_MACC_. In reality, *MACCT* transcript level may change over time, affecting the value of *k*_MACC_ (Martin & Saftner, 1995) suggesting *MACCT* levels are critical to correctly model MACC. To improve the model fit for MACC, a putative timecourse of *k*_MACC_ was estimated using a piecewise cubic Hermite polynomial with six knots (Fig. 8). This time dependent *k*_MACC_ now reflects its regulation through a time dependent *MACCT* expression. The predicted MACCT activity in winter fruit decrease throughout ripening, while in summer fruit it was predicted to peak at ORG stage (day 5). The observation for summer fruit aligns with findings in cv. ‘Ailsa Craig’ (Martin & Saftner, 1995) where the MACC activity sharply increased when fruit start to ripen, peak at ORG stage, and declined in ripe fruit.

**Figure 8.**
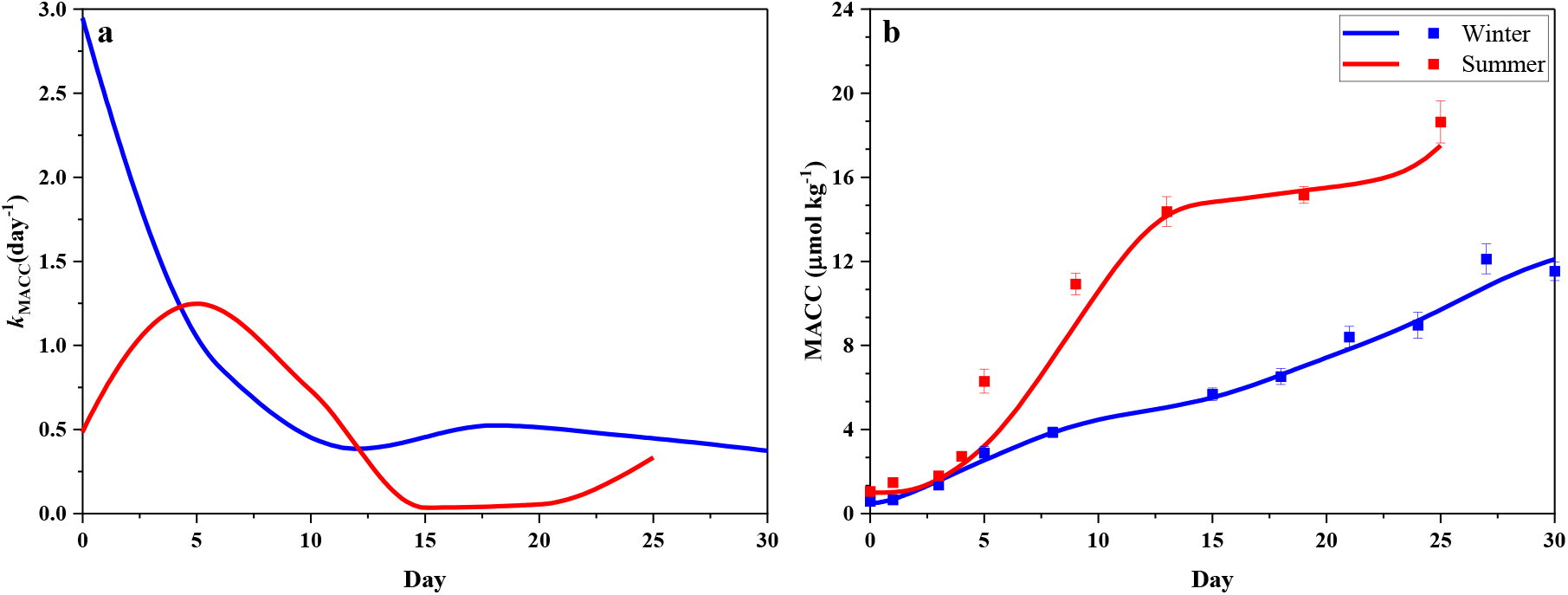
Model improvement of the prediction of MACC turnover winter and summer tomatoes during fruit ripening and postharvest storage. (a) *k*_MACC_ parameter, (b) MACC content. The lines are the model predictions, while the points are measurement data. Error bars represent the standard error of the mean (n = 5 for winter fruit and n = 4 for summer fruit).

The ethylene biosynthesis model from Van de Poel et al. (2014a) did not have experimental data on protein abundance and assumed ACO1 to be the main contributor to ACO activity. We were able to identify the contribution of each individual ACO member (ACO1, ACO5 and ACO6) to ethylene biosynthesis in both winter and summer tomatoes (Fig. 3). After introducing *f*_HKG_ to eliminate the difference in HKG expression, there was no seasonal difference in *k*_p,ACO5_ and *k*_p,ACO6_. However, *k*_p,ACO1_ and especially *k*_p,ACO3_ were lower in summer than in winter tomatoes (Table 2). Also, *k*_d,ACO_ was smaller in summer tomato compared to in winter fruit. This suggests that the target genes themselves were affected by the growing season and/or their coressponding proteins were differentially regulated.

The model results on ACOs and total ACS showed that the translation rate was much faster than the degradation rate, which was in agreement with previous studies recording the importance of trancription levels and translation in controling protein abundance during tomato deveopement and ripening (Hinkson & Elias, 2011; Belouah *et al*., 2019). However, at summer BR stage (day 3), transcription and translation could not explain the protein abundance of ACO5 and ACO6 (Fig. 3). This may suggest the presence of additional protein protection mechanism through, for instance, heat shock proteins (Vierling, 1991). The modeled enzyme activity of each ACO member revealed its contribution to the overall ACO activity and thus ethylene biosynthesis. Given the fixed *k*_ACO3_, the higher simulated protein abundance of ACO3 in winter compared to summer fruit suggests that ACO3 contributes more to ethylene production in winter than in summer fruit (Fig. 3). While ACO1 remains at a low basal level in winter fruit, it is the main isozyme contributing to ethylene production in summer fruit (Supplementary Fig. 3). This finding is somewhat surprising given other research showed that ACO1, predominantly regulates ethylene production (Barry *et al*., 1996; Xiong *et al*., 2005). An expression study of ACO1, ACO2, and ACO3 in transformed yeast showed that the ACO1 strain had the highest activity, while the ACO3 and ACO2 strains reached 65 % and 45 % of the maximum activity of ACO1 (Bidonde *et al*., 1998).

### Modeling ethylene signaling reveals the complex regulation of signaling proteins

The discrepancy between simulated and measured protein abundance of ETR2 and ETR3 could be due to regulations not incorporated in the model. Previous studies have shown that, ETR1 also interacts with RTE1, a positive regulator of ETR1 which stabilizes ETR1 and promotes its signaling (Zhou *et al*., 2007; Resnick *et al*., 2008; Qiu *et al*., 2012; Chang *et al*., 2014). On the other hand, TPR1, a tetratricopeptide repeat protein, has been found to interact with ETR1 and ETR3, inducing ethylene responses (Lin *et al*., 2008b, 2009a). However, it is unclear whether TPR1 (*i*) competes with CTRs for binding to ETRs, resulting in inactive CTRs or (*ii*) inactivates ETRs, inducing their degradation. These interactions of ETRs with other proteins indicate that the receptor turnover not just relies on its gene expression and the ubiquitin-related degradation pathway.

Based on small amino acid differences, ETRs could potentially show different ethylene-binding affinities (Azhar *et al*., 2023). O’Malley *et al*. (2005) showed that the ethylene-binding affinities of tomato receptors (ETR1, ETR2, ETR3) varied more than those in *Arabidopsis*. However, their analysis was semi-quantitative and only considered ethylene-bing domains of homodimers of ETR1, ETR2, and ETR3, ignoring any heterodimers. Therefore, this information was insufficient to use in our model. Acknowledging the limited data we still proceeded with their estimation to preliminarily examine the ethylene-binding affinities for homodimers and heterodimers (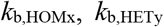; Table 3) suggesting different ethylene-binding affinities for some heterodimers. More detailed information on ethylene-binding affinity could help to better estimate those rate constants.

During signaling, EIN2 cleaves into a C-terminal (EIN2_Cterm_) and N-terminal part (EIN2_Nterm_) (Binder, 2020) (Fig. 1). EIN2_Cterm_ downregulates *EBF1*/2 mRNAs (Li *et al*., 2015) and translocates to the nucleus to activate ethylene responses via the EIN3/EIL family of transcription and ethylene response factors (Zhang *et al*., 2017). Since the experimental data on CTR1 and EIN2 is limited to the membrane bound fraction, the model leaves their abundance and signal transmission in the nucleus undetermined. Additional measurement quantifying subcellular translocation of EIN2_Cterm_ using LC/MS, might be helpful to estimate the parameter of EIN2 cleavage (Qiao *et al*., 2012). Previous studies showed that 1-MCP and its analogs reduced the expression of ethylene biosynthesis and signaling genes, indicating a feedback mechanism in transcription, influenced by ethylene signaling (Nakatsuka *et al*., 1998; Xu *et al*., 2016; Mata *et al*., 2021). In our model, this feedback mechanism was simplified into EIN2_Cterm_ directly regulating the transcription of ethylene response genes. We tried to use this feedback mechanism to describe the experimental data on ethylene responses (e.g. ethylene regulated gene expressions currently used as model inputs). However, this was not feasible due to a lack of quantitative information on EIN2_Cterm_ in terms of its total abundance, its turnover and translocation to the nucleus. In addition, no data on the specific contribution of EIN2_Cterm_ to specific ethylene response genes is available. There is also no information on possible post-transcriptional modifications. Therefore, modeling this feedback mechanism into the model is challenging and remains to be investigated.

### Model validation reveals different regulation during on- and off-vine ripening

The *in-vitro* ACS activity could not be well modeled (Fig. 5d, 6d, 7d). Maybe, different ACS members might become predominant under different conditions, like with ACO. Due to the lack of protein data on ACS, the regulation of ACS turnover, was lumped over the different ACS members, reflected by the generic parameters *k*_p,ACS_ and *k*_d,ACS_. Additionally, as the turnover is actually regulated by several mechanisms including translation, ubiquitination, phosphorylation, and dimerization (Park *et al*., 2021), these mechanisms might be differently regulated during on- and off-vine ripening, which was not fully captured by the model. This might also explain observed deviations for ACO activity (Fig. 5e, 6e, 7e). 1-MCP was predicted to delay the rise of the total ACS protein abundance, and to a lesser extent of the ACO members (Supplementary Fig. 8). These findings align with those of Tassoni (2006) who observed that 1-MCP caused a delayed increase in combined ACS2, ACS4, ACS6 transcription and protein levels, while it suppressed to a smaller extent the combined transcription and protein of ACO1, ACO3, ACO4 during fruit ripening. 1-MCP was predicted to strongly affect the abundance of most receptors (Supplementary Fig. 11), in agreement with the findings of Mata *et al*. (2021).

## Conclusions

An integrated kinetic model of the ethylene biosynthesis and signaling pathways was developed. The ethylene biosynthesis pathway under different conditions was largely dictated by the gene expression data, similar to what was observed by Van de Poel et al. (2014a). While the ethylene signaling model encompassed all interactions, including receptor dimer formation, dimer/CTR interaction, and dimer/CTR/EIN2 interaction, the experimental data were not rich enough to provide explicit information on all these different protein configurations. As a result, the involved model parameters stay relatively undefined. The analysis of protein complexes such as co-immunoprecipitation or immunoblotting in yeast two-hybrid systems is recommended to elucidate homo- and heteromeric interactions among receptors as well as the interaction of ETRs and CTRs. Utilizing ethylene-binding assay with transgenic yeast is a viable approach for investigating the binding capacity of receptors to ethylene. To improve the explanation of the model on ethylene signaling, the analysis of CTRs, EIN2, and downstream signaling proteins in nucleus is needed. The model structure of the ethylene signaling part is the premise for a further extensive model when information related to protein – protein interactions is sufficiently available. Additionally, experimental data is needed to extend the model with regard to the activation of specific ethylene response factors and the biochemical reactions triggered by these, thereby closing the loop of ethylene biosynthesis and signaling. The next level of this model should allow us to predict physiological responses such as respiration rate, and softening. Such a model would be instrumental to further understand climacteric ripening of tomato fruit.

## Acknowledgements

This research has been funded by the Vietnam National Foundation for Science and Technology Development (NAFOSTED) under grant number FWO.106-NN.2017.01, the VLIR-UOS Global Minds programme 2018, and KU Leuven (project C14/22/076).

## Author’s contributions

TMVN collected data, developed the kinetic model and simulations, and drafted the manuscript. DTT revised the manuscript. Clara I. Mata revised he manuscript. BVDP, BN, and MLATMH supervised the study, contributed to the model development and revised the manuscript. All authors read and approved the final manuscript.

## Data availability

The data that support the findings of this study are available from the corresponding author upon reasonable request.

