## Supplementary Dataset 1 for "Kinetic modeling of ethylene biosynthesis and signaling pathways in ripening tomato fruit"

```
%------------------------------------------------------------------------
%                              Ethylene biosynthesis and Signaling model
%------------------------------------------------------------------------
% The translation from mRNA to protein: The translation is assumed to be solely controlled by gene expression with a specific rate constant
% kp. The protein degradation is regulated by  a general degradation rate kd.
% mRNA -> p            with a specific rate constant kp
% p -> d               with a general rate constant kd
%
% Ethylene biosynthesis
% SAM converts to ACC which is catalyzed by the total of ACS2,4,6
% SAM + ACS2,4,6 -> ACC + ACS2,4,6                 with a general constant kacs
%
% ACC converts to C2H4 which is catalyzed by ACO1,3,5,6. ACC also converts to MACC.
% ACC + ACO1,3,5,6 -> C2H4 + ACO1,3,5,6 + MACC      with a rate constant kaco1,3,5,6 and  a rate constant kmacc
%
% C2H4 diffused (C2H4dif) to environment with a rate contant kdifC2H4
% C2H4 -> C2H4interal + C2H4 dif
%
% Ethylene signaling
% Receptors first form homodimer and heterodimer then interact with CTR1 (we assumed CTR1 is mainly contribute to the signal transmission).
% These two steps are simplied into one reaction
% pETR + pETR + CTR1 -> pETR_ETR_CTR1 with a rate constant kpCPX
%
% The degradation of homodimer and heterodimer are enhanced by C2H4 with kbhom and kbhet.
%
% The complex pETR-CTR faster the degration of EIN2.
% EIN2 > EIN2_C_term   with a rate constant kcterm. EIN2_C_term is the last product.
%
%------------------------------------------------------------------------
% ParamDef: Initialising parameters.
%-------------------------------------------------------------------------------------
<ParamDef>
      %-------------------------------------------------------------------------------------
      % To be compareable , the values are set as the estimate values. Orange represents winter and green (as comments) represents summer season.
      % Those values only in orange are estimated  in common or fixed.Intial values of ACC, MACC, C2H4, ACO1,5,6, receptors, CTR1 and EIN2 are
      % fixed based on the on-vine experimental results at IMG stage. ACO3 and ACS are arbitrarily fixed with in the range of the interpolation
      % of the rate constants of ACO1,3,6.
      % Note: those values are original, before coverting units.
      %-------------------------------------------------------------------------------------
      <Param> kpACO1 = 16355.5595; %8567.7908
      <Param> kpACO3 = 20.5152; %0.39454
      <Param> kpACO5 = 34.0673; %31.3055
      <Param> kpACO6 = 15.341; %23.3186
      <Param> kdACO = 0.78354; %0.26258
      <Param> kpACS = 94.1997; %174.2655
      <Param> kdACS = 0.1; %fixed
      <Param> kACS = 0.0001; %fixed
      <Param> kACO1 = 1.034e-08; %1.1567e-07
      <Param> kACO3 = 0.001; %fixed
      <Param> kACO5 = 0.0058986; %0.0012777
      <Param> kACO6 = 2.0659e-06; %0.00017648
      <Param> kdifC2H4 = 1.168;
      <Param> kMACC = 1.1354; %0.79603
      <Param> caco = 16.7402;
      <Param> cacs = 0.21199;
      <Param> fHKG = 1; %2.07
      <Param> SAM_0 = 14.8023; %35.8856
      <Param> a2 = 5.0223; %5.8328
      <Param> a3 = 1.4642; %4.3377
      <Param> a4 = 2.148; %3.3208
      <Param> a5 = 2.177; %2.3789
      <Param> a6 = 1.5522; %3.7755
      <Param> kpETR1 = 7003.5748; %45446.0389
      <Param> kpETR2 = 25.095; %25910.5244
      <Param> kpETR3 = 136.4589; %216.0504
      <Param> kpETR4 = 0.44736; %0.15707
      <Param> kpETR6 = 38.7377; %55.9444
      <Param> kpETR7 = 154.5573; %52.3263
      <Param> kdETR = 0.1; %fixed
      <Param> kpCTR1 = 0.90485; %1.9025
      <Param> kdCTR1 = 0; %fixed
      <Param> kpEIN2 = 2.3379; %6.0612
      <Param> kdEIN2 = 0.02; %fixed
      <Param> kpCPX = 10; %fixed
      <Param> kdCPX = 0; %fixed
      <Param> kbhom1 = 1.1504; %14.6361
      <Param> kbhet1 = 0.36416; %1774.3679
      <Param> kbhet3 = 0.26755; %0.0074877
      <Param> kbhom2 = 0.0073099; %0.0040965
      <Param> kbhet2 = 0.60444; %0.19314
      <Param> kbhet4 = 0.0021968; %6.7825e-05
```

```
     <Param> kbCPX = 0.1; %fixed
     <Param> ACC_0 = 0;
     <Param> MACC_0 = 0.5;%1
     <Param> C2H4_0 = 0.01;%0.04
     <Param> ACO1_0 = 500000; %500000
     <Param> ACO3_0 = 2;
     <Param> ACO5_0 = 30;%20
     <Param> ACO6_0 = 50;   %80
     <Param> ACS_0 = 100;%1.6
     <Param> ETR1_0 = 55;   %55
     <Param> ETR2_0 = 2;    %2
     <Param> ETR3_0 = 120;%150
     <Param> ETR4_0 = 0;
     <Param> ETR6_0 = 300;%380
     <Param> ETR7_0 = 600;%200
     <Param> CTR1_0 = 0;
     <Param> EIN2_0 = 6;%4.6
     <Param> ETR1_ETR1_CTR1_0 = 1.2;
     <Param> ETR2_ETR2_CTR1_0 = 1.2;
     <Param> ETR3_ETR3_CTR1_0 = 1.2;
     <Param> ETR1_ETR4_CTR1_0 = 0.45;
     <Param> ETR1_ETR6_CTR1_0 = 0.45;
     <Param> ETR1_ETR7_CTR1_0 = 0.45;
     <Param> ETR2_ETR4_CTR1_0 = 0.05;
     <Param> ETR2_ETR6_CTR1_0 = 0.05;
     <Param> ETR2_ETR7_CTR1_0 = 0.05;
     <Param> ETR3_ETR4_CTR1_0 = 1.2;
     <Param> ETR3_ETR6_CTR1_0 = 1.2;
     <Param> ETR3_ETR7_CTR1_0 = 1.2;
     <Param> ETR1_ETR2_CTR1_0 = 1.2;
     <Param> ETR1_ETR3_CTR1_0 = 1.2;
     <Param> ETR2_ETR3_CTR1_0 = 1.2;
     <Param> kcEIN2 = 0.1; %fixed

%-------------------------------------------------------------------------------------

</ParamDef>

%-------------------------------------------------------------------------------------
<StateDef>

     %-------------------------------------------------------------------------------------
     cTP = 12995720.4/10^9;% (µg TP/kg tissue) % cTP is the conversion factor from total protein (TP) to tissue
     cMP = 68482.62032/10^9;% (µg MP/kg tissue) % cMP is the conversion factor from total protein (TP) to tissue
     %-------------------------------------------------------------------------------------
     % Declaring and initialising ODE based state variables.
     %-------------------------------------------------------------------------------------
     <State> ACO1 = ACO1_0*cTP;
     <State> ACO3 = ACO3_0*cTP;
     <State> ACO5 = ACO5_0*cTP;
     <State> ACO6 = ACO6_0*cTP;
     <State> ACS = ACS_0*cTP;
     <State> ACC = ACC_0;
     <State> MACC = MACC_0;
     <State> C2H4 = C2H4_0;
     <State> ETR1 = ETR1_0*cMP;
     <State> ETR2 = ETR2_0*cMP;
     <State> ETR3 = ETR3_0*cMP;
     <State> ETR4 = ETR4_0*cMP;
     <State> ETR6 = ETR6_0*cMP;
     <State> ETR7 = ETR7_0*cMP;
     <State> CTR1 = CTR1_0*cMP;
     <State> EIN2 = EIN2_0*cMP;
     <State> ETR1_ETR1_CTR1 = ETR1_ETR1_CTR1_0*cMP;
     <State> ETR2_ETR2_CTR1 = ETR2_ETR2_CTR1_0*cMP;
     <State> ETR3_ETR3_CTR1 = ETR3_ETR3_CTR1_0*cMP;
     <State> ETR1_ETR4_CTR1 = ETR1_ETR4_CTR1_0*cMP;
     <State> ETR1_ETR6_CTR1 = ETR1_ETR6_CTR1_0*cMP;
     <State> ETR1_ETR7_CTR1 = ETR1_ETR7_CTR1_0*cMP;
     <State> ETR2_ETR4_CTR1 = ETR2_ETR4_CTR1_0*cMP;
     <State> ETR2_ETR6_CTR1 = ETR2_ETR6_CTR1_0*cMP;
     <State> ETR2_ETR7_CTR1 = ETR2_ETR7_CTR1_0*cMP;
     <State> ETR3_ETR4_CTR1 = ETR3_ETR4_CTR1_0*cMP;
     <State> ETR3_ETR6_CTR1 = ETR3_ETR6_CTR1_0*cMP;
     <State> ETR3_ETR7_CTR1 = ETR3_ETR7_CTR1_0*cMP;
     <State> ETR1_ETR2_CTR1 = ETR1_ETR2_CTR1_0*cMP;
     <State> ETR1_ETR3_CTR1 = ETR1_ETR3_CTR1_0*cMP;
     <State> ETR2_ETR3_CTR1 = ETR2_ETR3_CTR1_0*cMP;
     %-------------------------------------------------------------------------------------
</StateDef>
```

```
%------------------------------------------------------------------------------
% ModelDef: Defining derivatives of the state variables.
%------------------------------------------------------------------------------
<ModelDef>
      %------------------------------------------------------------------------------
      cTP = 12995720.4/10^9;
      cMP = 68482.62032/10^9;
      % Total ETR_ETR_CTR1 complex:
      pETR_CTR_tot = ETR1_ETR1_CTR1+ETR2_ETR2_CTR1+ETR3_ETR3_CTR1+ETR1_ETR4_CTR1+ETR1_ETR6_CTR1+ETR1_ETR7_CTR1+ETR2_ETR4_CTR1
                   +ETR2_ETR6_CTR1+ETR2_ETR7_CTR1+ETR3_ETR4_CTR1+ETR3_ETR6_CTR1+ETR3_ETR7_CTR1+ETR1_ETR2_CTR1+ETR1_ETR3_CTR1
                   +ETR2_ETR3_CTR1;
      % A putative SAM concentration is contructed specifically for winter and summer season. trange is the total experimental time o f winter/summer
      % season
      trange=max(t_cond);
      SAM_t=max(0,pchip([0:trange/5:trange],[SAM_0 a2 a3 a4 a5 a6],t));

      %------------------------------------------------------------------------------
      % Defining derivatives.
      %------------------------------------------------------------------------------
      %ACO, assume the translation rate is specific and the degradation rate is general
      Deriv(ACO1)= kpACO1*cTP*fHKG*mRNA_ACO1(t)-kdACO*ACO1;
      Deriv(ACO3)= kpACO3*cTP*fHKG*mRNA_ACO3(t)-kdACO*ACO3;
      Deriv(ACO5)= kpACO5*cTP*fHKG*mRNA_ACO5(t)-kdACO*ACO5;
      Deriv(ACO6)= kpACO6*cTP*fHKG*mRNA_ACO6(t)-kdACO*ACO6;
      %ACS, assume the translation and degradation rates are general
      Deriv(ACS)= fHKG*kpACS*cTP*(mRNA_ACS2(t)+mRNA_ACS4(t)+mRNA_ACS6(t))-kdACS/(1+C2H4)*ACS;
      %ACC, MACC and C2H4 turnovers
      Deriv(ACC) = (kacs/cTP)*SAM_t*ACS+(-kaco1*ACC*ACO1-kaco3*ACC*ACO3-kaco5*ACC*ACO5-kaco6*ACC*ACO6)/cTP-kmacc*ACC;
      Deriv(MACC)= kmacc*ACC;
      Deriv(C2H4)= (kaco1*ACC*ACO1+kaco3*ACC*ACO3+kaco5*ACC*ACO5+kaco6*ACC*ACO6)/cTP-kdifC2H4*C2H4;
      % ETR turnover of Subfamily I
      Deriv(ETR1) = kpETR1*cMP*fHKG*mRNA_ETR1(t)-kdETR*ETR1-kpCPX/cMP^2*CTR1*(2*ETR1^2+ETR1*(ETR2+ETR3)+ETR1*(ETR4+ETR6+ETR7))...
                    -ETR1*C2H4*(kbhom_1/cMP*2*ETR1+kbhet1/cMP*ETR2+kbhet2/cMP*ETR3+kbhet3/cMP*(ETR4+ETR6+ETR7));
      Deriv(ETR2) = kpETR2*cMP*fHKG*mRNA_ETR2(t)-kdETR*ETR2-kpCPX/cMP^2*CTR1*(2*ETR2^2+ETR2*(ETR1+ETR3)+ETR2*(ETR4+ETR6+ETR7))...
                    -kbhom_1/cMP*C2H4*2*ETR2*ETR2-kbhet1/cMP*C2H4*ETR2*ETR1...
                    -kbhet2/cMP*C2H4*ETR2*ETR3-kbhet3/cMP*C2H4*ETR2*(ETR4+ETR6+ETR7);
      Deriv(ETR3) = kpETR3*cMP*fHKG*mRNA_ETR3(t)-kdETR*ETR3-kpCPX/cMP^2*CTR1*(2*ETR3^2+ETR3*(ETR1+ETR2)+ETR3*(ETR4+ETR6+ETR7))...
                    -kbhom2/cMP*C2H4*2*ETR3*ETR3...
                    -kbhet2/cMP*C2H4*ETR3*(ETR1+ETR2)-kbhet4/cMP*C2H4*ETR3*(ETR4+ETR6+ETR7);
      % ETR turnover of Subfamily II
      Deriv(ETR4) = kpETR4*cMP*fHKG*mRNA_ETR4(t)-kdETR*ETR4-kpCPX/cMP^2*ETR4*CTR1*(ETR1+ETR2+ETR3)...
                    -kbhet3/cMP*C2H4*ETR4*(ETR1+ETR2)...
                    -kbhet4/cMP*C2H4*ETR4*ETR3;
      Deriv(ETR6) = kpETR6*cMP*fHKG*mRNA_ETR6(t)-kdETR*ETR6-kpCPX/cMP^2*ETR6*CTR1*(ETR1+ETR2+ETR3)...
                    -kbhet3/cMP*C2H4*ETR6*(ETR1+ETR2)...
                    -kbhet4*C2H4*ETR6*ETR3;
      Deriv(ETR7) = kpETR7*cMP*fHKG*mRNA_ETR7(t)-kdETR*ETR7-kpCPX/cMP^2*ETR7*CTR1*(ETR1+ETR2+ETR3)...
                    -kbhet3/cMP*C2H4*ETR7*(ETR1+ETR2)...
                    -kbhet4/cMP*C2H4*ETR7*ETR3;
      % CTR1 turnover
      Deriv(CTR1) = kpCTR1*cMP*fHKG*mRNA_CTR1(t)-kdCTR1*CTR1-kpCPX/cMP^2*CTR1*(ETR1^2+ETR2^2+ETR3^2)...
                    -kpCPX/cMP^2*CTR1*(ETR1*ETR2+ETR1*ETR3+ETR2*ETR3)...
                    -kpCPX/cMP^2*CTR1*(ETR1*(ETR4+ETR6+ETR7)+ETR2*(ETR4+ETR6+ETR7)+ETR3*(ETR4+ETR6+ETR7));
      % Homodier_CTR1 turnover
      Deriv(ETR1_ETR1_CTR1) = kpCPX/cMP^2*ETR1^2*CTR1-kbCPX*C2H4*ETR1_ETR1_CTR1;
      Deriv(ETR2_ETR2_CTR1) = kpCPX/cMP^2*ETR2^2*CTR1-kbCPX*C2H4*ETR2_ETR2_CTR1;
      Deriv(ETR3_ETR3_CTR1) = kpCPX/cMP^2*ETR3^2*CTR1-kbCPX*C2H4*ETR3_ETR3_CTR1;
      % Heterodimer_CTR1 turnover
      Deriv(ETR1_ETR2_CTR1) = kpCPX/cMP^2*CTR1*ETR1*ETR2-kdCPX*ETR1_ETR2_CTR1-kbCPX*C2H4*ETR1_ETR2_CTR1;
      Deriv(ETR1_ETR3_CTR1) = kpCPX/cMP^2*CTR1*ETR1*ETR3-kdCPX*ETR1_ETR3_CTR1-kbCPX*C2H4*ETR1_ETR3_CTR1;
      Deriv(ETR1_ETR4_CTR1) = kpCPX/cMP^2*CTR1*ETR1*ETR4-kdCPX*ETR1_ETR4_CTR1-kbCPX*C2H4*ETR1_ETR4_CTR1;
      Deriv(ETR1_ETR6_CTR1) = kpCPX/cMP^2*CTR1*ETR1*ETR6-kdCPX*ETR1_ETR6_CTR1-kbCPX*C2H4*ETR1_ETR6_CTR1;
      Deriv(ETR1_ETR7_CTR1) = kpCPX/cMP^2*CTR1*ETR1*ETR7-kdCPX*ETR1_ETR7_CTR1-kbCPX*C2H4*ETR1_ETR7_CTR1;
      Deriv(ETR2_ETR3_CTR1) = kpCPX/cMP^2*CTR1*ETR2*ETR3-kdCPX*ETR2_ETR3_CTR1-kbCPX*C2H4*ETR2_ETR3_CTR1;
      Deriv(ETR2_ETR4_CTR1) = kpCPX/cMP^2*CTR1*ETR2*ETR4-kdCPX*ETR2_ETR4_CTR1-kbCPX*C2H4*ETR2_ETR4_CTR1;
      Deriv(ETR2_ETR6_CTR1) = kpCPX/cMP^2*CTR1*ETR2*ETR6-kdCPX*ETR2_ETR6_CTR1-kbCPX*C2H4*ETR2_ETR6_CTR1;
      Deriv(ETR2_ETR7_CTR1) = kpCPX/cMP^2*CTR1*ETR2*ETR7-kdCPX*ETR2_ETR7_CTR1-kbCPX*C2H4*ETR2_ETR7_CTR1;
      Deriv(ETR3_ETR4_CTR1) = kpCPX/cMP^2*CTR1*ETR3*ETR4-kdCPX*ETR3_ETR4_CTR1-kbCPX*C2H4*ETR3_ETR4_CTR1;
      Deriv(ETR3_ETR6_CTR1) = kpCPX/cMP^2*CTR1*ETR3*ETR6-kdCPX*ETR3_ETR6_CTR1-kbCPX*C2H4*ETR3_ETR6_CTR1;
      Deriv(ETR3_ETR7_CTR1) = kpCPX/cMP^2*CTR1*ETR3*ETR7-kdCPX*ETR3_ETR7_CTR1-kbCPX*C2H4*ETR3_ETR7_CTR1;
      % EIN2 turnover
      Deriv(EIN2) = kpEIN2*cMP*fHKG*mRNA_EIN2(t)-(kdEIN2/cMP*pETR_CTR_tot)*EIN2-kcEIN2*EIN2;
      %------------------------------------------------------------------------------
</ModelDef>

%------------------------------------------------------------------------------
% TransOut: Processing ODE model outputs.
%------------------------------------------------------------------------------
<TransOut>
      %------------------------------------------------------------------------------
      trange=max(t_cond);
```

```
    cTP = 12995720.4/10^9;
    cMP = 68482.62032/10^9;

    %------------------------------------------------------------------------------------
    % Assigning transformed values to existing ODE based state variables.
    %------------------------------------------------------------------------------------
    pETR_CTR_tot = ETR1_ETR1_CTR1+ETR2_ETR2_CTR1+ETR3_ETR3_CTR1+ETR1_ETR4_CTR1+ETR1_ETR6_CTR1+ETR1_ETR7_CTR1+ETR2_ETR4_CTR1
    +ETR2_ETR6_CTR1+ETR2_ETR7_CTR1+ETR3_ETR4_CTR1+ETR3_ETR6_CTR1+ETR3_ETR7_CTR1+ETR1_ETR2_CTR1+ETR1_ETR3_CTR1+ETR2_ETR3_CTR1) /cMP;
    %------------------------------------------------------------------------------------
    % Declaring and assigning values to new output variables.
    %------------------------------------------------------------------------------------
    <Output> C2H4prod = kdifC2H4*24*C2H4/24;
    <Output> ETR1_out = (ETR1+2*ETR1_ETR1_CTR1+ETR1_ETR4_CTR1+ETR1_ETR6_CTR1+ETR1_ETR7_CTR1+ETR1_ETR2_CTR1+ETR1_ETR3_CTR1)/cMP;
    <Output> ETR2_out = (ETR2+2*ETR2_ETR2_CTR1+ETR2_ETR4_CTR1+ETR2_ETR6_CTR1+ETR2_ETR7_CTR1+ETR1_ETR2_CTR1+ETR2_ETR3_CTR1)/cMP;
    <Output> ETR3_out = (ETR3+2*ETR3_ETR3_CTR1+ETR3_ETR4_CTR1+ETR3_ETR6_CTR1+ETR3_ETR7_CTR1+ETR1_ETR3_CTR1+ETR2_ETR3_CTR1)/cMP;
    <Output> ETR4_out = (ETR4+ETR1_ETR4_CTR1+ETR2_ETR4_CTR1+ETR3_ETR4_CTR1)/cMP;
    <Output> ETR6_out = (ETR6+ETR1_ETR6_CTR1+ETR2_ETR6_CTR1+ETR3_ETR6_CTR1)/cMP;
    <Output> ETR7_out = (ETR7+ETR1_ETR7_CTR1+ETR2_ETR7_CTR1+ETR3_ETR7_CTR1)/cMP;
    <Output> CTR1_out = (CTR1+ETR1_ETR1_CTR1+ETR1_ETR4_CTR1+ETR1_ETR6_CTR1+ETR1_ETR7_CTR1+ETR2_ETR2_CTR1+ETR2_ETR4_CTR1
           +ETR2_ETR6_CTR1+ETR2_ETR7_CTR1   +ETR3_ETR3_CTR1+ETR3_ETR4_CTR1+ETR3_ETR6_CTR1+ETR3_ETR7_CTR1 +ETR1_ETR2_CTR1
           +ETR1_ETR3_CTR1+ETR2_ETR3_CTR1)/cMP;
    <Output> ACOact_out = (kaco1*ACO1+kaco3*ACO3+kaco5*ACO5+kaco6*ACO6)*caco*24/cTP/24;
    <Output> ACSact_out = kacs*ACS*cacs*24/cTP/24;
    <Output> SAM_out = max(0,pchip([0:trange/5:trange],[SAM_0 a2 a3 a4 a5 a6 ],t_mod));
    <Output> ACO1_out = ACO1/cTP;
    <Output> ACO5_out = ACO5/cTP;
    <Output> ACO6_out = ACO6/cTP;
    <Output> EIN2_out = EIN2/cMP;
    <Output> ACS_out = ACS/cTP;
    %------------------------------------------------------------------------------------
</TransOut>
```
