## Supplementary Figures for "Kinetic modeling of ethylene biosynthesis and signaling pathways in ripening tomato fruit"

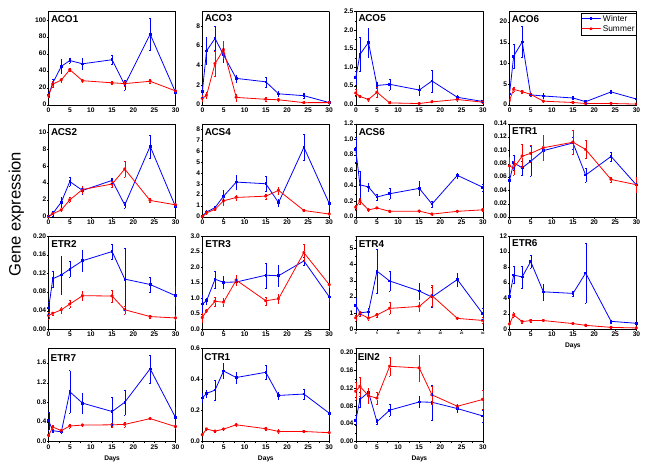

Supplementary Fig. 1. Gene expression of winter and summer tomatoes during on-vine ripening. The *f*_HKG_ is included in gene expression in summer fruit.

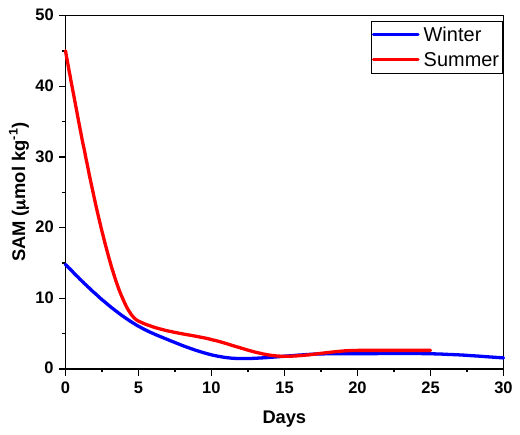

Supplementary Fig. 2. The estimation of SAM concentration during on-vine ripening

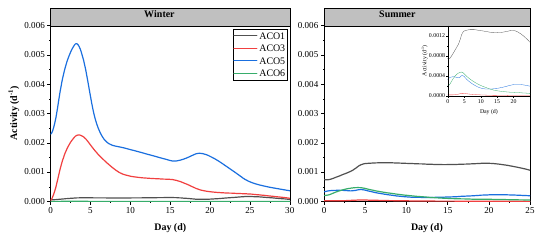

Supplementary Fig. 3. The modeled activity of ACO1, 3, 5, 6 in winter and summer fruit during on-vine ripening.

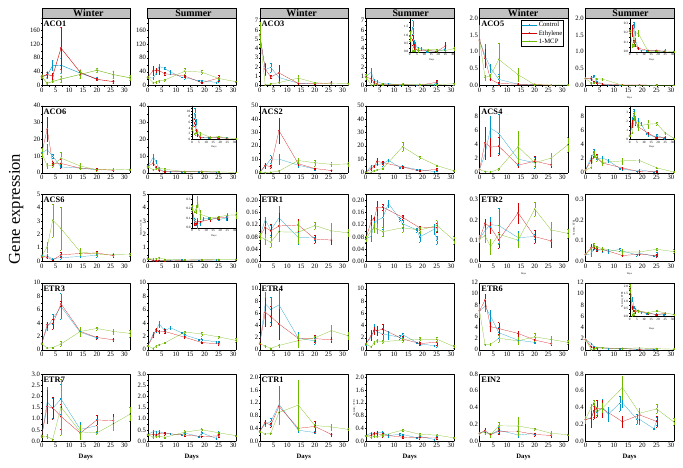

Supplementary Fig. 4. Gene expression of during off-vine postharvest ripening. The *f*_HKG_ is included in gene expression in summer fruit.

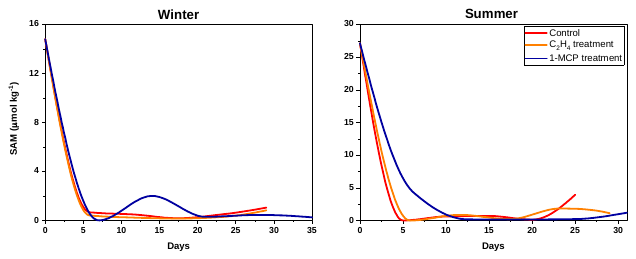

Supplementary Fig. 5. The estimation of SAM concentration during off-vine postharvest ripening

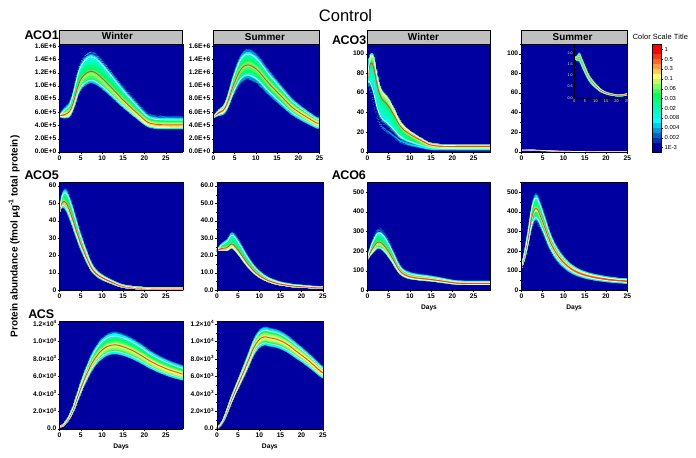

Supplementary Fig. 6. The prediction of the abundance of ethylene biosynthesis proteins of control winter and summer tomatoes during off-vine postharvest ripening.

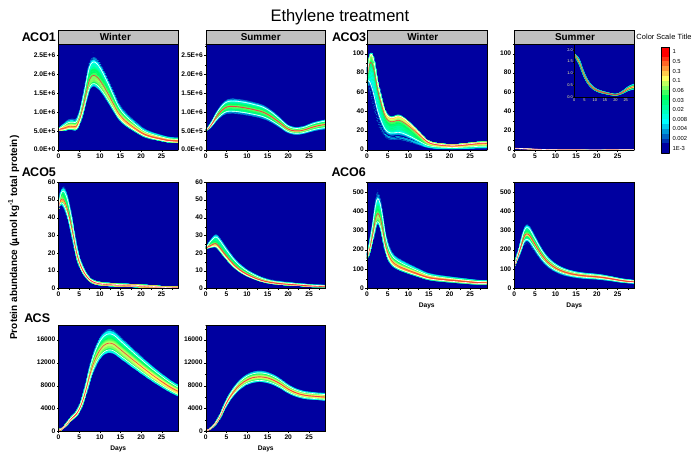

Supplementary Fig. 7. The prediction of the abundance of ethylene biosynthesis proteins of ethylene treated winter and summer tomatoes during off-vine postharvest ripening.

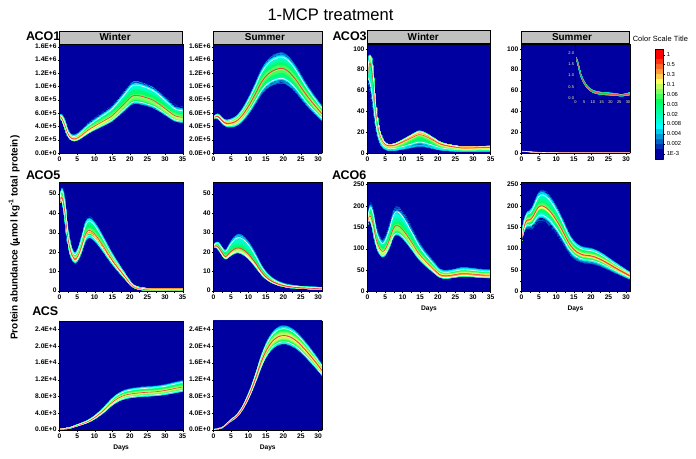

Supplementary Fig. 8. The prediction of the abundance of ethylene biosynthesis proteins of 1-MCP treated winter and summer tomatoes during off-vine postharvest ripening.

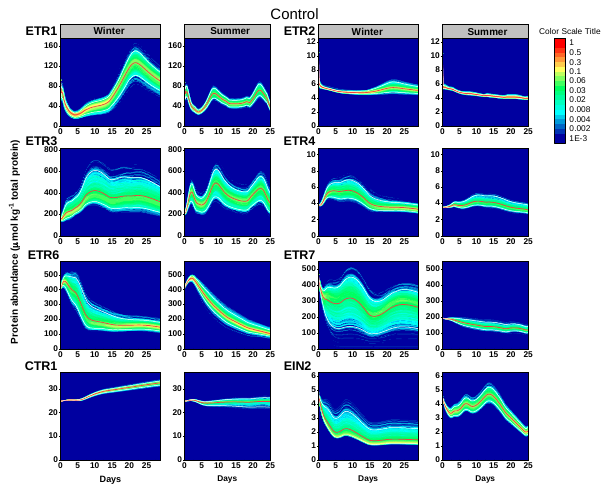

Supplementary Fig. 9. The prediction of the abundance of ethylene signaling proteins of control winter and summer tomatoes during off-vine postharvest ripening.

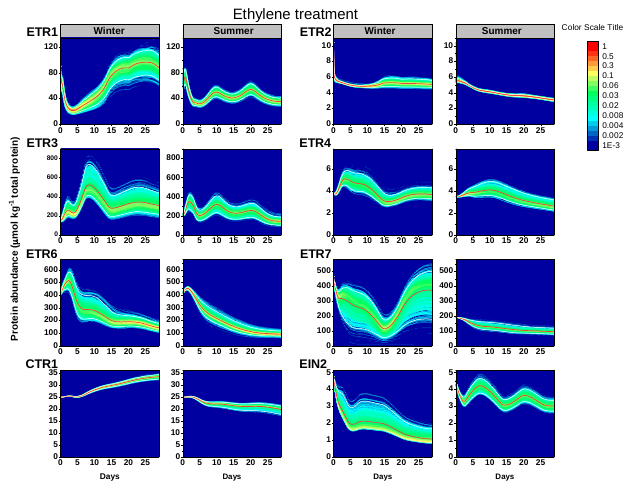

Supplementary Fig. 10. The prediction of the abundance of ethylene signaling proteins of ethylene treated winter and summer tomatoes during off-vine postharvest ripening.

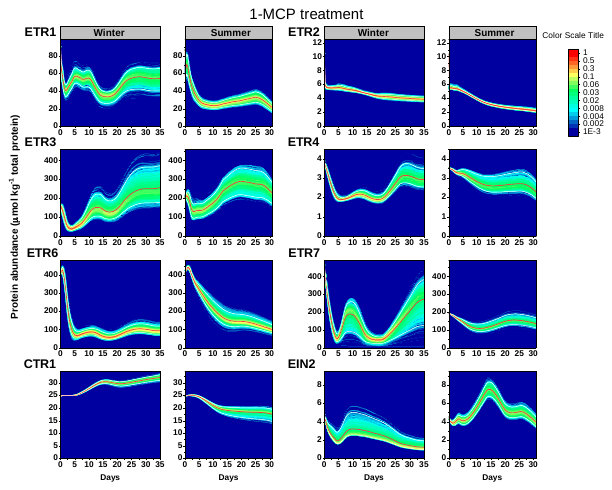

Supplementary Fig. 11. The prediction of the abundance of ethylene signaling proteins of 1-MCP treated winter and summer tomatoes during off-vine postharvest ripening.
