## Supplementary Tables for "Kinetic modeling of ethylene biosynthesis and signaling pathways in ripening tomato fruit"

Supplementary Table 1. The time interval according to winter and summer control fruit.

| Stage | Storage time (days) | |
| --- | --- | --- |
|  | **Winter_Control** | **Summer_Control** |
| IMG | 0 | 0 |
| MG | 1 | 1 |
| BR | 3 | 3 |
| TRN | 5 | 4 |
| ORG | 8 | 5 |
| LR | NA | 9 |
| RR | 15 | 13 |
| RR+3 | 18 | NA |
| RR+6 | 21 | 19 |
| RR+9 | 24 | NA |
| RR+12 | 27 | 25 |
| RR+15 | 30 | NA |

Supplementary Table 2. The description of rates, factors and compounds

| **Parameter** | **Description** (the rate of…) |
| --- | --- |
| 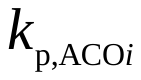 | specific production of ACO member *i* (translation) |
| 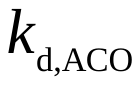 | general degradation of all ACO*i* members |
| 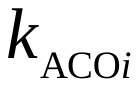 | specific ACO activity of member *i* |
| 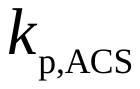 | general production of all ACS*i* members (translation) |
| 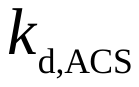 | general degradation of all ACS*i* members |
| 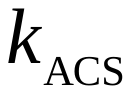 | general ACS activity |
| 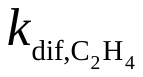 | ethylene diffusion |
| 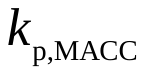 | production of MACC |
| 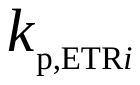 | specific production of ETR member *i* of subfamily I (translation) |
| 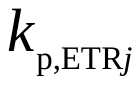 | specific production of ETR member *j* of subfamily II (translation) |
| 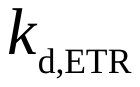 | general degradation of ETR member *i* and *j* of subfamily I and II |
| 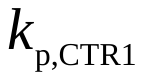 | production of CTR1 (translation) |
| 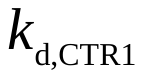 | degradation of CTR1 |
| 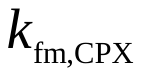 | production of 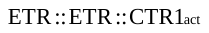 complex |
| 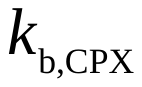 | ethylene binding to 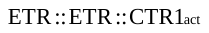 complex |
| 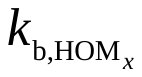 | ethylene binding to the various homodimers |
| 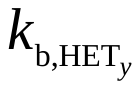 | ethylene binding to the various heterodimers |
| 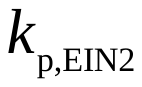 | production of EIN2 (translation) |
| 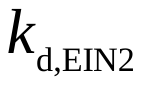 | degradation of EIN2 |
| 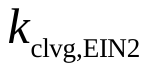 | cleavage of EIN2 |
| **Factor** |  |
| 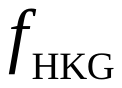 | housekeeping gene factor |
| 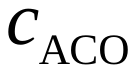 | conversion of ACO from *in vivo* to *in vitro* |
| 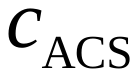 | conversion of ACS from *in vivo* to *in vitro* |
| **Compound** |  |
| 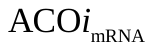 | ACO mRNA member *i* |
| 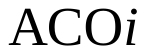 | ACO member *i* |
| 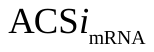 | ACS mRNA member *i* |
| 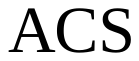 | all ACS members |
| 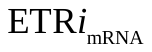 | ETR mRNA member *i* of subfamily I |
| 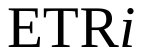 | ETR member *i* of subfamily I |
| 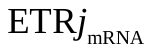 | ETR mRNA member *j* of subfamily II |
| 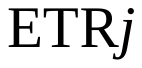 | ETR member *j* of subfamily II |
| 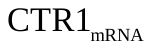 | CTR1 mRNA |
| 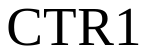 | CTR1 |
| 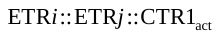  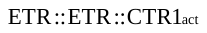  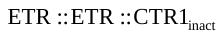 | active specific complex  active complex in general  inactivate complex |
| 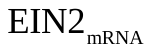 | EIN2 mRNA |
| 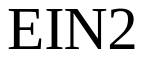 | EIN2 |
| EIN2_Cterm_ | EIN2 C terminal |
| EIN2_Nterm_ | EIN2 N terminal |
| SAM | SAM |
| ACC | ACC |
| MACC | MACC |
| C_2_H_4_ | C_2_H_4_ |

Supplementary Table 3. The initial values of ethylene biosynthesis, signaling in model calibration

| Parameter estimates | Winter tomato | Summer tomato | Unit |
| --- | --- | --- | --- |
| ACC_0 | 0 | 0 | µmol kg^-1^ ^¤^ |
| MACC_0 | 0.5 | 1 | µmol kg^-1^ ^¤^ |
| C2H4_0 | 0.01 | 0.04 | µmol kg^-1^ ^¤^ |
| pACO1_0 | 500000 | 500000 | µmol kg^-1^ * |
| pACO3_0 | 2 | 2 | µmol kg^-1^ * |
| pACO5_0 | 30 | 20 | µmol kg^-1^ * |
| pACO6_0 | 50 | 90 | µmol kg^-1^ * |
| pACS_0 | 100 | 1.6 | µmol kg^-1^ * |
| pETR1_0 | 55 | 1 | µmol kg^-1 #^ |
| pETR3_0 | 120 | 150 | µmol kg^-1 #^ |
| pETR4_0 | 0 | 0 | µmol kg^-1 #^ |
| pETR6_0 | 300 | 380 | µmol kg^-1 #^ |
| pETR7_0 | 600 | 200 | µmol kg^-1 #^ |
| pCTR1_0 | 0 | 0 | µmol kg^-1 #^ |
| pEIN2_0 | 6 | 4.7 | µmol kg^-1 #^ |
| pETR1_ETR1_CTR1_0 | 1.2 | 1.2 | µmol kg^-1 #^ |
| pETR2_ETR2_CTR1_0 | 1.2 | 1.2 | µmol kg^-1 #^ |
| pETR3_ETR3_CTR1_0 | 1.2 | 1.2 | µmol kg^-1 #^ |
| pETR1_ETR4_CTR1_0 | 0.45 | 0.45 | µmol kg^-1 #^ |
| pETR1_ETR6_CTR1_0 | 0.45 | 0.45 | µmol kg^-1 #^ |
| pETR1_ETR7_CTR1_0 | 0.45 | 0.45 | µmol kg^-1 #^ |
| pETR2_ETR4_CTR1_0 | 0.05 | 0.05 | µmol kg^-1 #^ |
| pETR2_ETR6_CTR1_0 | 0.05 | 0.05 | µmol kg^-1 #^ |
| pETR2_ETR7_CTR1_0 | 0.05 | 0.05 | µmol kg^-1 #^ |
| pETR3_ETR4_CTR1_0 | 1.2 | 1.2 | µmol kg^-1 #^ |
| pETR3_ETR6_CTR1_0 | 1.2 | 1.2 | µmol kg^-1 #^ |
| pETR3_ETR7_CTR1_0 | 1.2 | 1.2 | µmol kg^-1 #^ |
| pETR1_ETR2_CTR1_0 | 1.2 | 1.2 | µmol kg^-1 #^ |
| pETR1_ETR3_CTR1_0 | 1.2 | 1.2 | µmol kg^-1 #^ |
| pETR2_ETR3_CTR1_0 | 1.2 | 1.2 | µmol kg^-1 #^ |

^¤^ Expressed at a fresh weight base, ^*^ Expressed at a total protein base, ^#^ Expressed at the membrane protein base

Supplementary Table 4. The initial values of ethylene biosynthesis, signaling in model validation

| Parameter estimates | Winter tomato | | | Summer tomato | | | | | | | Unit |
| --- | --- | --- | --- | --- | --- | --- | --- | --- | --- | --- | --- |
|  | **Control** | | **1-MCP** | | **Ethylene** | | **Control** | **1-MCP** | **Ethylene** | |  |
| kmacc | 0.35 ±0.04 | | 0.078 ±0.007 | | 0.25 ±0.04 | | 0.45 ±0.003 | 0.091 ±0.000 | 0.25 ±0.025 | | d^-1^ |
| a2 | 0.68 ±0.18 | | 0.014 ±0.005 | | 0.43 ±0.23 | | 0.017 ±0.000 | 4.3 ±0.4 | 0.002 ±0.005 | | µmol kg^-1 ¤^ |
| a3 | 0.50 ±0.08 | | 2 .0±0.1 | | 0.24 ±0.11 | | 0.64 ±0.01 | 0.18 ±0.14 | 0.88 ±0.14 | | µmol kg^-1 ¤^ |
| a4 | 0.19 ±0.21 | | 0.31 ±0.13 | | 0.16 ±0.13 | | 0.71 ±0.01 | 0.16 ±0.06 | 0.22 ±0.19 | | µmol kg^-1 ¤^ |
| a5 | 0.51 ±0.28 | | 0.46 ±0.14 | | 0.29 ±0.20 | | 0.14 ±0.72 | 0.22 ±0.13 | 1.8 ±0.8 | | µmol kg^-1 ¤^ |
| a6 | 1.1 ±0.7 | | 0.26 ±0.37 | | 0.84 ±0.70 | | 4.0 ±1.3 | 1.2 ±0.4 | 1.1 ±1.3 | | µmol kg^-1 ¤^ |
| Parameters fixed as common for three treatments per season | | | | | | | | | | | |
| SAM_0 | 14.802 | | | | | 27.13 | | | | | µmol kg^-1 ¤^ |
| ACC_0 | 0 | | | | | 0 | | | | | µmol kg^-1 ¤^ |
| MACC_0 | 0.65909 | | | | | 1.479 | | | | | µmol kg^-1 ¤^ |
| C2H4_0 | 0.034 | | | | | 0.046 | | | | | µmol kg^-1 ¤^ |
| pACO1 | 5.51E+05 | | | | | 5.20E+05 | | | | | µmol kg^-1^ * |
| pACO5_0 | 45.5 | | | | | 22.762 | | | | | µmol kg^-1^ * |
| pACO6_0 | 158.9 | | | | | 119.74 | | | | | µmol kg^-1^ * |
| pACS_0 | 210 | | | | | 97.637 | | | | | µmol kg^-1^ * |
| pETR1_0 | 84.5 | | | | | 80.402 | | | | | µmol kg^-1 #^ |
| pETR2_0 | 5.51 | | | | | 5.4006 | | | | | µmol kg^-1 #^ |
| pETR3_0 | 155 | | | | | 197.39 | | | | | µmol kg^-1 #^ |
| pETR4_0 | 2.17 | | | | | 1.8218 | | | | | µmol kg^-1 #^ |
| pETR6_0 | 406 | | | | | 409 | | | | | µmol kg^-1 #^ |
| pETR7_0 | 466 | | | | | 191.2 | | | | | µmol kg^-1 #^ |
| pCTR1_0 | 12.5 | | | | | 12.517 | | | | | µmol kg^-1 #^ |
| pEIN2_0 | 4.8 | | | | | 4.5259 | | | | | µmol kg^-1 #^ |
| Parameters fixed as common for winter and summer season | | | | | | | | | | | |
| pETR1_ETR1_CTR1_0 | | 1.2 | | | | | | | | µmol kg^-1 #^ | |
| pETR2_ETR2_CTR1_0 | | 1.2 | | | | | | | | µmol kg^-1 #^ | |
| pETR3_ETR3_CTR1_0 | | 1.2 | | | | | | | | µmol kg^-1 #^ | |
| pETR1_ETR4_CTR1_0 | | 0.45 | | | | | | | | µmol kg^-1 #^ | |
| pETR1_ETR6_CTR1_0 | | 0.45 | | | | | | | | µmol kg^-1 #^ | |
| pETR1_ETR7_CTR1_0 | | 0.45 | | | | | | | | µmol kg^-1 #^ | |
| pETR2_ETR4_CTR1_0 | | 0.05 | | | | | | | | µmol kg^-1 #^ | |
| pETR2_ETR6_CTR1_0 | | 0.05 | | | | | | | | µmol kg^-1 #^ | |
| pETR2_ETR7_CTR1_0 | | 0.05 | | | | | | | | µmol kg^-1 #^ | |
| pETR3_ETR4_CTR1_0 | | 1.2 | | | | | | | | µmol kg^-1 #^ | |
| pETR3_ETR6_CTR1_0 | | 1.2 | | | | | | | | µmol kg^-1 #^ | |
| pETR3_ETR7_CTR1_0 | | 1.2 | | | | | | | | µmol kg^-1 #^ | |
| pETR1_ETR2_CTR1_0 | | 1.2 | | | | | | | | µmol kg^-1 #^ | |
| pETR1_ETR3_CTR1_0 | | 1.2 | | | | | | | | µmol kg^-1 #^ | |
| pETR2_ETR3_CTR1_0 | | 1.2 | | | | | | | | µmol kg^-1 #^ | |

^*^ Expressed at a total protein base, ^#^ Expressed at the membrane protein base
